# Cxcr4 desensitization is an essential regulatory mechanism controlling the extra-follicular B cell response

**DOI:** 10.1101/537761

**Authors:** Nagham Alouche, Amélie Bonaud, Vincent Rondeau, Julie Nguyen, Etienne Cricks, Niclas Setterblad, Matthieu Mahevas, Karl Balabanian, Marion Espéli

## Abstract

The signaling axis formed by the chemokine CXCL12 and its receptor CXCR4 plays an important role in B cell development and activation and is finely regulated by a process termed desensitization. Mutations leading to a truncation of the C-terminus tail of CXCR4 and thus to a defective desensitization have been reported in two diseases, a rare immunodeficiency called the WHIM syndrome and a B cell plasmacytoma called Waldenstrom’s Macroglobulinemia (WM). How CXCR4 desensitization may impact B cell activation in the context of a T-independent extra-follicular response is still unknown. Here using a unique mouse model bearing an orthologous gain of function mutation of *Cxcr4* we report that Cxcr4 desensitization is an essential gatekeeper controlling B lymphocyte entry into cycle, plasma cell differentiation, migration and maturation upon Myd88-dependent signaling. Altogether, our results support an essential role for Cxcr4 desensitization in limiting the depth and width of the B cell extra-follicular response and PC development.

## Introduction

The signalling induced by the G protein-coupled receptor Cxcr4 upon binding of its ligand the chemokine Cxcl12 plays an essential role in B cell homeostasis. Indeed, Cxcl12, although known as SDF-1, was originally identified as an early B cell survival factor^1,2^. Since then numerous studies have reinforced the relevance of this signalling axis for B cell development. Several works have notably established its central role for the regulation of the germinal centre (GC) reaction and for plasma cell (PC) homing to and persistence within the BM^3–8^. However, the role of this axis in the extra-follicular humoral response has never been formally addressed.

Cxcr4/Cxcl12 signalling is tightly regulated by a mechanism called desensitization that relies on the phosphorylation of the C-terminus tail of the receptor and the subsequent recruitment of β-arrestins. This mechanism is absolutely required to uncouple the G-proteins from the receptor and thus for shutting down the Cxcl12-induced signalling. Recruitment of the β-arrestins then leads to receptor internalization through clathrin-coated pits and to its degradation or recycling to the surface^9^. Our group and others showed that this desensitization process is defective in a rare immunodeficiency called the WHIM syndrome (WS)^10,11^. Most WS patients present mutations in the C-ter tail of CXCR4 thus abrogating the desensitization of the receptor and leading to a gain of function characterized by a hyperactive receptor upon CXCL12 binding^10,12^. To decipher the cellular and molecular mechanisms underlying the physiopathology of the WS we generated a knock-in mouse model bearing one of the mutations identified in WS patients (mutation at the 1013 position and leading to a 15 amino acid truncation)^13^. These mice referred to as *Cxcr4*^+/*1013*^, phenocopied most of the WS features including a gain of function of Cxcr4 upon Cxcl12 stimulation and a severe pan-leukopenia^13^. Moreover, using this unique model we recently showed that the gain of function of Cxcr4 affects in a B-cell intrinsic manner the T-dependent immune response and recapitulates the vaccination failures reported in WS patients. In addition, we observed an early aberrant accumulation of immature plasmablasts (PBs) in the BM following immunization in absence of Cxcr4 desensitization^7^.

Strikingly, the same (C1013G) mutation of *CXCR4* has also been reported in Waldenström’s Macroglobulinemia (WM), a rare indolent lymphoplasmacytic B-cell lymphoma characterized by an IgM peak and BM infiltration of malignant B cells with a phenotype going from activated B cells to PCs^14,15^. This somatic *CXCR4* mutation is observed in 20-30% of WM patients, is always found in conjunction with a *MYD88* gain of function mutation (itself presents in 90% of patients) and is associated with poor prognosis, BM involvement and therapy resistance^16,17^. How these two mutations synergize is still unclear but, together with our recent findings^7^, this raises the question of the impact of Cxcr4 desensitization in the context of MyD88 signalling on B cell and PC biology. We thus took advantage of our unique mouse model to unravel the implication of Cxcr4 desensitization on these processes. Our results demonstrate both *in vitro* and *in vivo* that Cxcr4 desensitization is an essential mechanism that dampens MyD88-mediated PC differentiation, maturation and migration. Importantly, we also report that Cxcr4 desensitization is a regulatory mechanism that controls B cell entry into cycle. Altogether our results demonstrate that Cxcr4 signalling needs to be tightly regulated to limit the extent of the B cell extra-follicular response.

## Results

### Cxcr4 desensitization limits Myd88-mediated plasma cell differentiation *in vitro* and *in vivo*

To assess how the Cxcr4 gain of function resulting from the defective desensitization of the receptor may impact Myd88-mediated B cell activation we stimulated splenic B cells from WT and *Cxer4^+/1013^* mice with the TLR4 and TLR9 ligands, LPS and CpG, *in vitro* in presence or in absence of Cxcl12. As expected we observed a strong induction of PB differentiation 4 days after LPS and CpG stimulation in the WT conditions (Figure 1A-B and Figure S1A). In presence of the gain-of-function mutation of *Cxcr4*, PB differentiation induced by both TLR ligands was significantly enhanced compared to WT B cells (Figure 1A-B and Figure S1A). This increased differentiation translated into increased IgM concentration measured in the supernatant of mutant B cell cultures compared to the WT ones (Figure 1B). The addition of Cxcl12 in the stimulation mix led to a small, albeit not significant, increase of PB differentiation in both the WT and the *Cxcr4*^+/*1013*^ cultures compared to the condition without Cxcl12 (Figure 1B and Figure S1A). Interestingly, stimulation with Cxcl12 alone promoted some PB differentiation of *Cxcr4*^+/*1013*^ B cells, a phenomenon barely detectable in WT cultures (Figure 1B). This Cxcl12-mediated PC differentiation was abolished by addition of the Cxcr4 antagonist AMD-3100 thus confirming that Cxcr4 desensitization regulates this phenomenon (Figure S1B). Of note, this Cxcl12-mediated PC differentiation was not associated with as much proliferation as was observed with the LPS conditions (Figure S1C) and this limited expansion also translated in a smaller amount of IgM secreted in this condition compared to the conditions with LPS (Figure 1B right panel).

**Figure 1:**
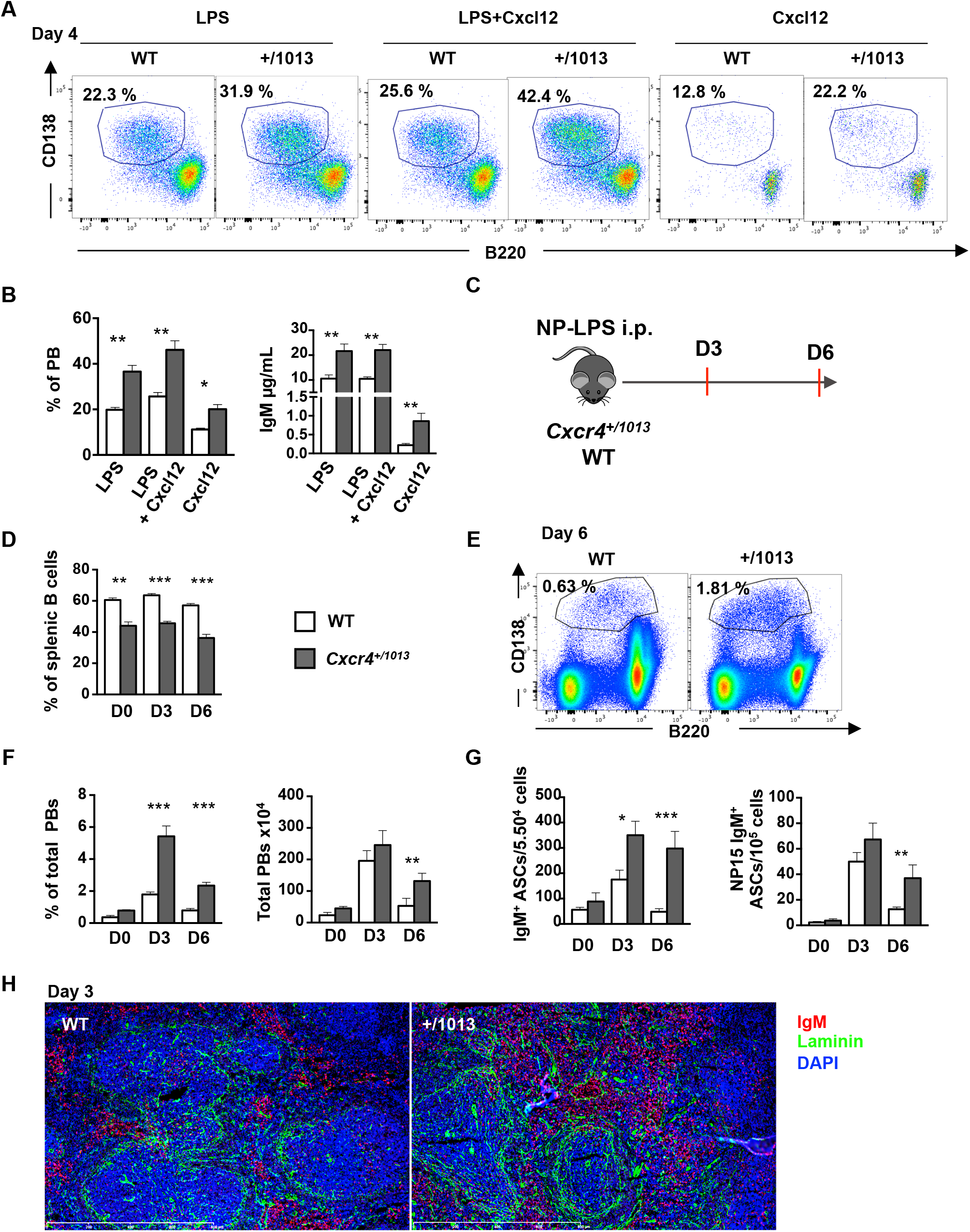
Cxcr4 desensitization limits Myd88-mediated plasma cell differentiation *in vitro* and *in vivo*: **(A-B)** Splenic B cells were cultured in presence of LPS, LPS+Cxcl12, or Cxcl12 alone for 4 days. **(A)** Representative dot plots for the gating of PBs (B220^lo^CD138^+^) generated in vitro after 4 days of culture. **(B)** The proportion of PBs was determined by FACS (left panel). IgM concentrations in the supernatants were determined by ELISA (right panel). **(C)** Schematic diagram for the NP-LPS immunisation. **(D)** Frequency of splenic B cells (CD19^+^ B220^+^) during NP-LPS immunisation. **(E)** Representative dot plots for PC gating in the spleen of both WT and *Cxcr4*^+/*1013*^ mice 6 days post NP-LPS immunisation. **(F)** Percentage and total number of PBs in the spleen at day 0, 3 and 6 post immunization. **(G)** Quantification of total IgM^+^ and NP15-specific IgM^+^ ASCs in the spleen by ELISpot at day 0, 3 and 6 post immunization. **(H)** Representative staining of spleen sections from WT and *Cxcr4*^+/*1013*^ mice at day 3 post immunization. PBs are stained with an anti-IgM Ab (red), basal membrane is stained with an anti-laminin Ab (green) and nuclei are stained with Hoechst 33342 (Blue), Scale bare: 800μm. **(I)** Heatmap showing the relative expression of different mRNA from WT and *Cxcr4*^+/*1013*^ splenic PBs determined by Biomark multiplex qPCRs at day 6 post immunization. The heatmap was generated using the heatmapper column Z score based on (2-^Δ^Ct^^) values. **(J)** Relative expression of Blimp1/GFP MFI in splenic PCs from *Cxcr4*^+/*1013*^ mice compared to the WT ones. Results are from 3 independent experiments (A-H and J) or one representative experiment out of 2 (I) (Mean ± SEM, n=3-4). Mann–Whitney U test was used to assess statistical significance (*P<0,05, ** P<0,01, ***P<0,001).

We next determined *in vivo* whether the gain of function of Cxcr4 was impacting the TLR-mediated Type 1 T-independent B cell immune response. WT and *Cxcr4*^+/*1013*^ mice were immunized i.p. with NP-LPS and the splenic immune response was evaluated at day 3 and 6 post injection (Figure 1C). The immunization was not correcting the B cell lymphopenia observed in the spleen of *Cxcr4*^+/*1013*^ mice^13^ (Figure 1D). This was mostly due to the reduced frequency and number of follicular B cells in mutant mice (Figure S1D). As previously reported, the frequency of marginal zone (MZ) B cells was increased and their number was normal in *Cxcr4*^+/*1013*^ mice compared to WT mice^13^ and this was not altered by NP-LPS immunisation (Figure S1D).

We further analysed splenic plasmablast (PB) generation and observed a significant increase of the frequency of PBs from day 3 that was still prominent at day 6 in *Cxcr4*^+/*1013*^ mice compared to WT ones (Figure 1E-F). Accordingly, we detected an increased frequency of total IgM^+^ antibody-secreting cells (ASCs) by ELISpot in the spleen of *Cxcr4*^+/*1013*^ mice compared to WT mice (Figure 1G, left panel), and an increased serum titre of total IgM in the mice bearing the Cxcr4 gain of function (Figure S1E, left panel). Moreover, the frequencies of NP-specific IgM^+^ ASCs were increased at day 6 in presence of the Cxcr4 gain of function (Figure 1G right and Figure S1E right). Importantly, the T-independent type I immunisation was not only enhanced but also more sustained in *Cxcr4*^+/*1013*^ mice compared to WT mice as demonstrated by the still elevated frequency and number of PBs observed at day 6 in the mutant mice while these numbers were back to baseline in the WT animals (Figure 1F-G). Analysis of splenic PBs by immunohistofluorescence revealed an increased detection in sections and also a more diffuse distribution in the red pulp in *Cxcr4*^+/*1013*^ spleens compared to WT spleens (Figure 1H).

Altogether, these results suggest that Cxcr4 desensitization is an important regulator of PC differentiation.

### Cxcr4 desensitization limits the entry of LPS-stimulated B cells into cycle

We addressed the cellular mechanism underlying this enhanced PB generation in absence of Cxcr4 desensitization *in vitro*. We first showed that the gain of function of Cxcr4 was not affecting B cell and PB survival by performing Annexin V/DAPI and active Caspase 3 staining on LPS-mediated *in vitro* differentiation of WT and mutant splenocytes at day 2 and 4 (Figure S2A). This was supported by the analysis of the expression of key apoptosis regulators in PBs from WT and *Cxcr4*^+/*1013*^ mice at day 4 after LPS stimulation (Figure S2B). We then evaluated whether proliferation may be impacted by the gain of function of Cxcr4. Splenic B cells were loaded with CTV and stimulated with LPS in absence or presence of Cxcl12. The CTV dilution was assessed from day 2 and up to day 4. Strikingly, we observed at day 2 that more mutant B cells had proliferated compared to WT B cells (Figure 2A-B). However, this difference was not detected anymore from day 3 of culture (Figure 2B). The addition of Cxcl12 to the culture did not significantly increase B cell proliferation compared to the LPS-only condition but we still observed more proliferation at day 2 in B cells mutant for *Cxcr4* (Figure S2C). Cxcl12 on its own was only inducing minimal B cell proliferation however, mutant B cells were cycling more than their WT counterparts (Figure S1C and S2C). We also measured CTV dilution in newly generated PBs and did not observe difference in the proliferation of WT and mutant PBs at any time point analysed in presence or in absence of Cxcl12 (Figure 2C-D and Figure S2D). Of note, in presence of Cxcl12 only we observed more proliferation for the mutant PBs compared to the WT ones at day 2 but this difference was not detectable anymore the following days (Figure S2D). These results brought us to consider whether it was not the proliferation of B cells that was affected but rather their entry in cycle. We thus, measured the frequency of B cells in G0, G1, S and G2-M phases following LPS stimulation. From day 1 and up to day 3 we observed a significant increase in the percentage of B cells in cycle (G1, S and G2-M phases) in *Cxcr4*^+/*1013*^ cultures compared to WT ones (Figure 2E-F and data not shown for day 1). The addition of Cxcl12 to the stimulation medium did not significantly increase the cycling of B cells compared to the condition with LPS alone (Figure S2E). As expected, most PBs were cycling^18^ and no difference was observed between our two experimental groups (Figure 2F, right panel). Moreover, a transcriptomic analysis confirmed that mutant splenic B cells 6 days after LPS injection present a characteristic cycling signature that is not observed in WT B cells (Figure S2F). Previous reports demonstrated that B cell differentiation into PB necessitates at least 3 rounds of cell division^19^. We thus assessed whether this enhanced cycling of B cells may account for an accelerated differentiation. We cultured WT x Blimp1-GFP and *Cxcr4*^+/*1013*^ x Blimp-1 GFP B cells with LPS and assessed the kinetic of appearance of GFP by videomicroscopy. As shown in Figure 2G, some GFP+ cells were already detectable in *Cxcr4*^+/*1013*^ cultures as early as 46 hours post LPS stimulation while they started appearing in WT cultures almost 20 hours later (Figure 2G-H). Our results demonstrate that Cxcr4 desensitization is an essential regulator of splenic B cell entry into cycle and this enhanced cycling seems to translate into exacerbated PB differentiation.

**Figure 2:**
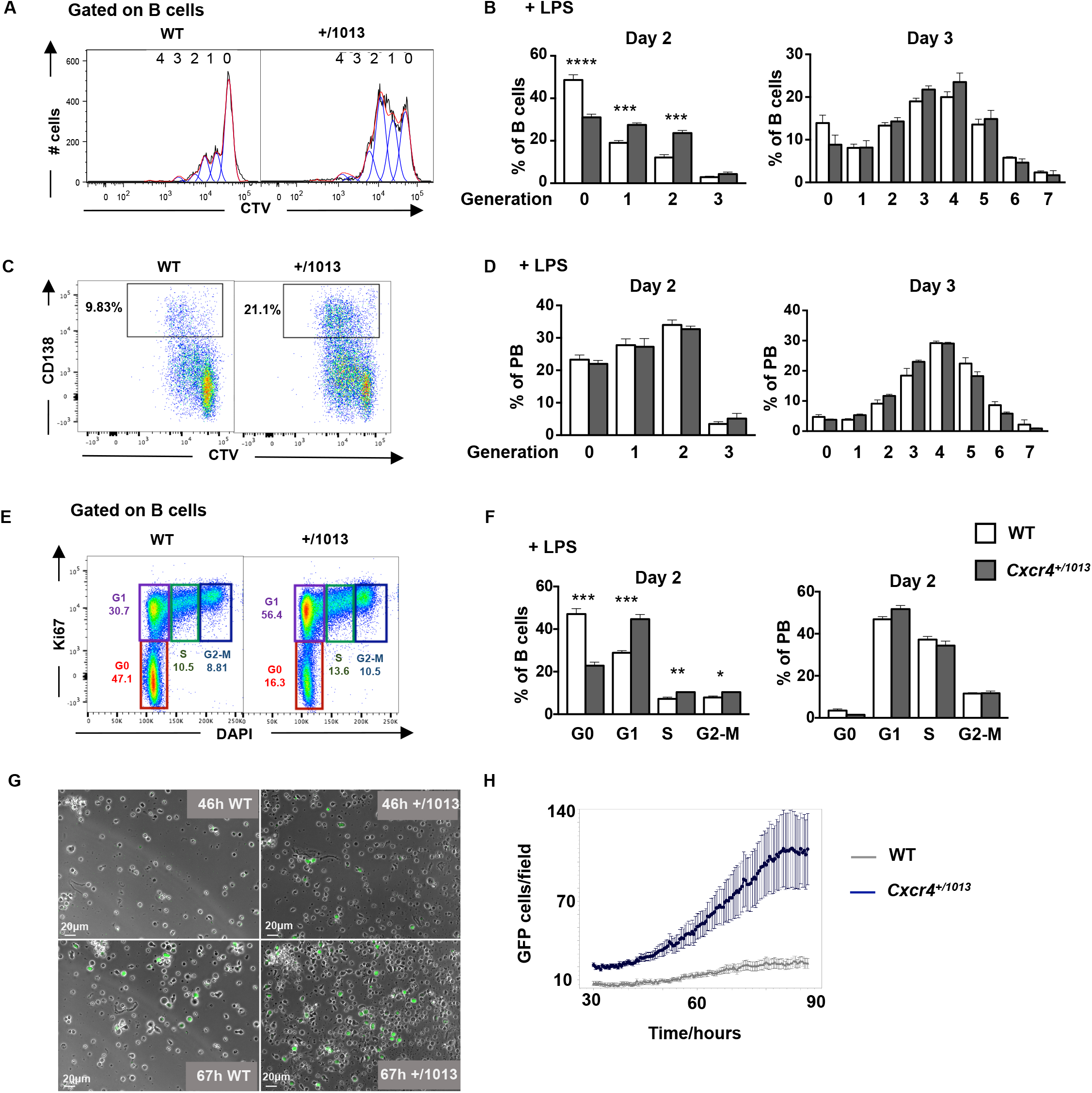
Cxcr4 desensitization inhibits the entry of LPS-stimulated B cells into cycle: **(A-D)** Splenocytes from WT and *Cxcr4*^+/*1013*^ mice were loaded with CTV and cultured in presence of LPS for 3 days. **(A)** Representative histograms of CTV dilution in LPS stimulated B cells at day 2. **(B)** Frequency of B cells present in each CTV-dilution generation at day 2 and 3. **(C)** Representative dot plots for CTV dilution during PBs (CD138^+^) differentiation at day 2 post LPS stimulation. **(D)** Frequency of PBs present in each CTV-dilution generation at day 2 and 3. **(E)** Representative plots showing the different phases of the cell cycle (G0, G1, S, G2-M) for splenic B cells at day 2 post LPS stimulation stained with DAPI and for Ki67. **(F)** Frequency of splenic B cells and PBs in each cell cycle phase at day 2 post LPS stimulation. **(G)** Representative fields from time-lapse imaging of splenic B cells from Blimp1^GFP/+^ x WT and Blimp1^GFP/+^ x *Cxcr4*^+/*1013*^ mice cultured in presence of LPS at 46 and 67h. Phase contrast and GFP fluorescence detection are shown. Scale bare: 20μm **(H)** Quantification of GFP+ cells/ imaged fields during LPS stimulation of splenic B cells. Results are from 2 independent experiments (A-F) or one representative experiment out of 2 (G-H) (Mean ± SEM, n=2-4). Two-tailed Student’s t tests was used to assess statistical significance (*P<0,05, ** P<0,01, ***P<0,001, ****P<0,0001).

### Cxcr4 desensitization controls follicular and MZ B cell differentiation into PB via distinct mechanisms

As shown previously^13^, *Cxcr4*^+/*1013*^ mice have reduced number of follicular B cells but normal number of MZ B cells. We thus wondered if the difference observed in terms of MyD88-mediated PB differentiation in absence of Cxcr4 desensitization could be explained by the different representation of these two subsets between WT and mutant mice. We thus sorted follicular and MZ B cells and differentiated them *in vitro* with LPS. We observed that both subsets were generating more PBs when they were bearing the *Cxcr4* gain of function mutation (Figure 3A). The analysis of the cell cycle at day 2 post-LPS stimulation showed that follicular B cells bearing the *Cxcr4* gain of function mutation were more in cycle than the WT ones (Figure 3B) with the frequency of cycling cells increasing from day 1 to day 3 (Figure 3B and Figure S3A (for day1)). As expected, after LPS stimulation MZ B cells were more in cycle than follicular B cells (Figure 3D-E)^20^. However, there was no significant difference in terms of cycle between mutant and WT MZ B cells at all time points analysed (Figure 3D-E and Figure S3B). To support these findings, we fed WT and *Cxcr4*^+/*1013*^ mice with EdU for 6 days and analysed EdU incorporation at day 6 and progressive EdU loss during a chase in splenic follicular and MZ B cells *in vivo* (Figure 3F). As shown in Figure 3G, more mutant follicular B cells had incorporated EdU compared to WT cells. In constrast, the frequency of EdU+, hence of cycling cells, was equivalent between WT and mutant MZ B cells at day 6, thus supporting our *in vitro* results. In addition, both mutant and WT follicular and MZ B cells lost the EdU labelling globally at the same rate during the chase suggesting that Cxcr4 desensitization does not impact their proliferation rate (Figure 3G). These results confirmed our *in vitro* results and suggest that the increased differentiation of mutant MZ B cells cannot be explained only by an enhanced cell cycle entry. We thus analysed more precisely the transcriptomic profile of MZ B cells in *Cxcr4*^+/*1013*^ mice and observed that their expression of the PC marker *Prdm1* was increased while the expression of the B cell marker *Bach2* was decreased (Figure 3H) suggesting that mutant MZ B cells display a transcriptional signature reminiscent of early PC differentiation. These differences were not detectable in follicular B cells (Figure S3C). Moreover, the percentage of Blimp1/GFP+ MZ B cells was increased in the mutant mice compared to the WT ones (Figure 3I and S3D). Altogether our results suggest that in absence of Cxcr4 desensitization both follicular and MZ B cells have an enhanced capacity to differentiate into PBs in response to LPS. In the case of follicular B cells this enhanced differentiation potential correlates with enhanced entry into cell cycle while in MZ B cells it is associated with an expression of transcription factors reminiscent of early PC differentiation.

**Figure 3:**
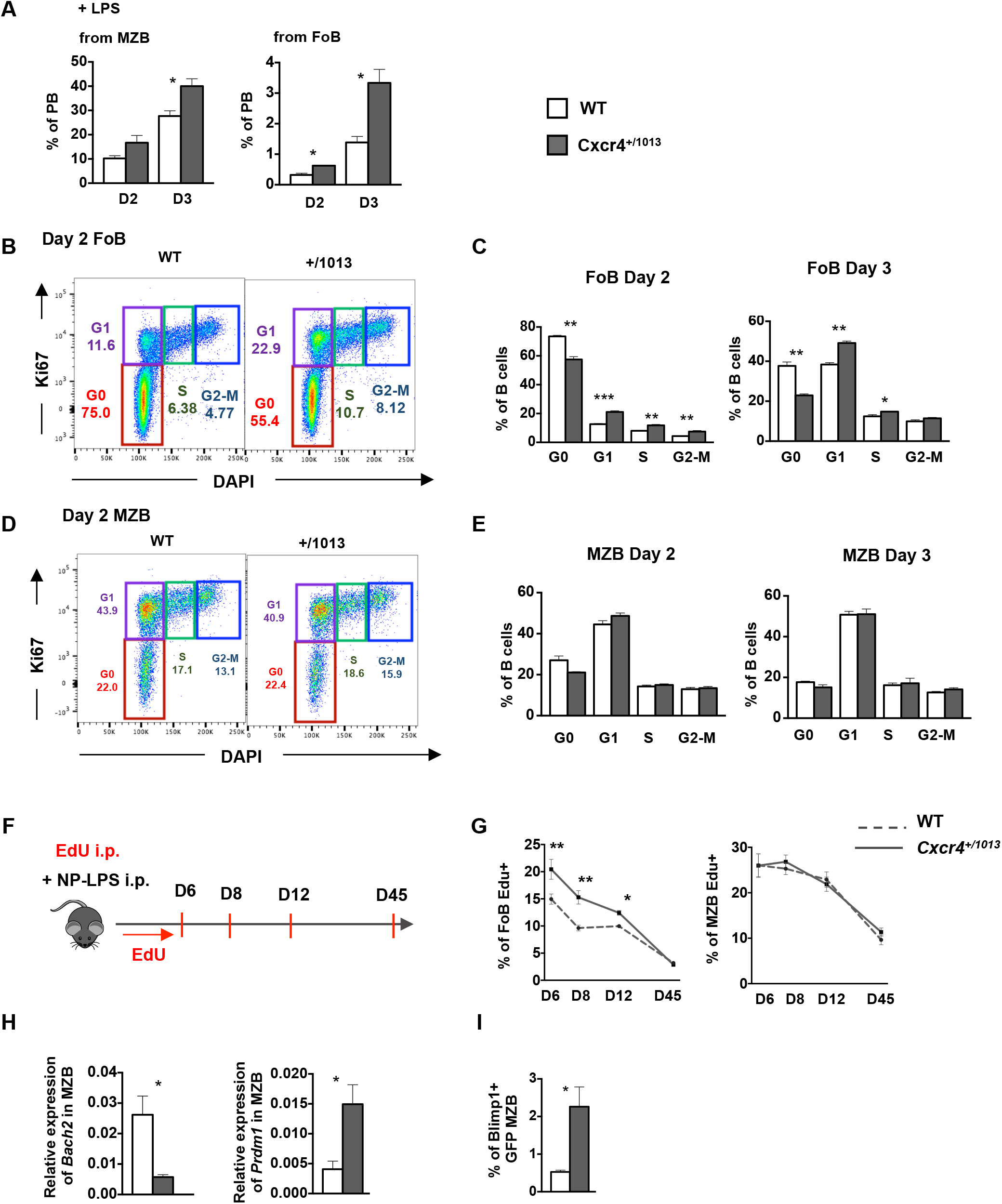
Cxcr4 desensitization controls follicular and MZ B cell differentiation into PB via distinct mechanisms: **(A-E)** MZB and FoB cells were sorted from WT and *Cxcr4*^+/*1013*^ mice spleens and stimulated in presence of LPS for 3 days. **(A)** Frequency of PBs generated from both MZ and Fo B cells after 2 and 3 days of LPS stimulation. **(B)** Representative dot plots of FoB cell cycle (G0, G1, S, G2-M) at day 2 post LPS stimulation detected with DAPI and Ki67 staining. **(C)** Frequency of FoB cells in each cell cycle phase at day 2 and 3 post LPS stimulation. **(D)** Representative dot plots of MZB cell cycle (G0, G1, S, G2-M) at day 2 post LPS stimulation detected with DAPI and Ki67 staining. **(E)** Frequency of MZB cells in each cell cycle phase at day 2 and 3 post LPS stimulation. **(F)** Schematic diagram for Edu labelling-chase and NP-LPS immunisation. **(G)** Frequency of EdU+ FoB and MZB cells from d6 and during the chase at the indicated time-points after NP-LPS immunisation. **(H)** Relative expression of *Bach2* and *Prdm1* in MZB cells presented as (2-^Δ^Ct^^). **(I)** Percentage of Blimp1-GFP+ cells amongst MZB cells at day 6 post NP-LPS immunisation. Results are from one representative experiment out of 2 (Mean ± SEM, n=2-5). Two-tailed Student’s t tests was used to assess statistical significance (*P<0,05, **P<0,01, ***P<0,001).

### Cxcr4 desensitization limits plasmablasts accumulation within the bone marrow

We previously reported that during a T-dependent response the desensitization of Cxcr4 was essential to limit the apparition of aberrant PBs within the BM presumably generated independently of the germinal centre response^7^. We thus investigated if such was the case in a humoral response that was independent of T cell help and not leading to GC formation. WT and *Cxcr4*^+/*1013*^ mice crossed to the Blimp1 GFP reporter strain were immunized i.p. with NP-LPS and their BM was analysed after 3 and 6 days (Figure 4A). As previously reported the BM cellularity was equivalent between WT and *Cxcr4*^+/*1013*^ mice^13^ (Figure 4B). We observed an increased detection of total IgM^+^ ASCs from day 3 and of NP-specific IgM^+^ ASCs from day 6 in the BM of *Cxcr4*^+/*1013*^ mice compared to the WT ones (Figure 4C-D). When we analysed the phenotype of BM PCs we observed, using the expression of the GFP as a readout of *Prdm1* expression and thus of PC differentiation, that immature PBs (GFP/Blimp-1^low^) were massively increased in the BM of mutant mice compared to WT ones as early as 3 days post immunisation (Figure 4E). Moreover, fully differentiated PCs (GFP/Blimp-1^high^) were significantly increased from day 6 in the BM of the *Cxcr4*^+/*1013*^ mice compared to their WT littermates (Figure 4E). To confirm these results, we assessed the number of total PCs in the BM of WT and mutant mice by immunohistofluorescence of thick BM sections. In support of our flow cytometry and ELISpot results, we enumerated 5 times more PCs per μm^2^ at day 6 after NP-LPS immunization in the BM sections of *Cxcr4*^+/*1013*^ mice compared to the WT ones (Figure 4F).

**Figure 4:**
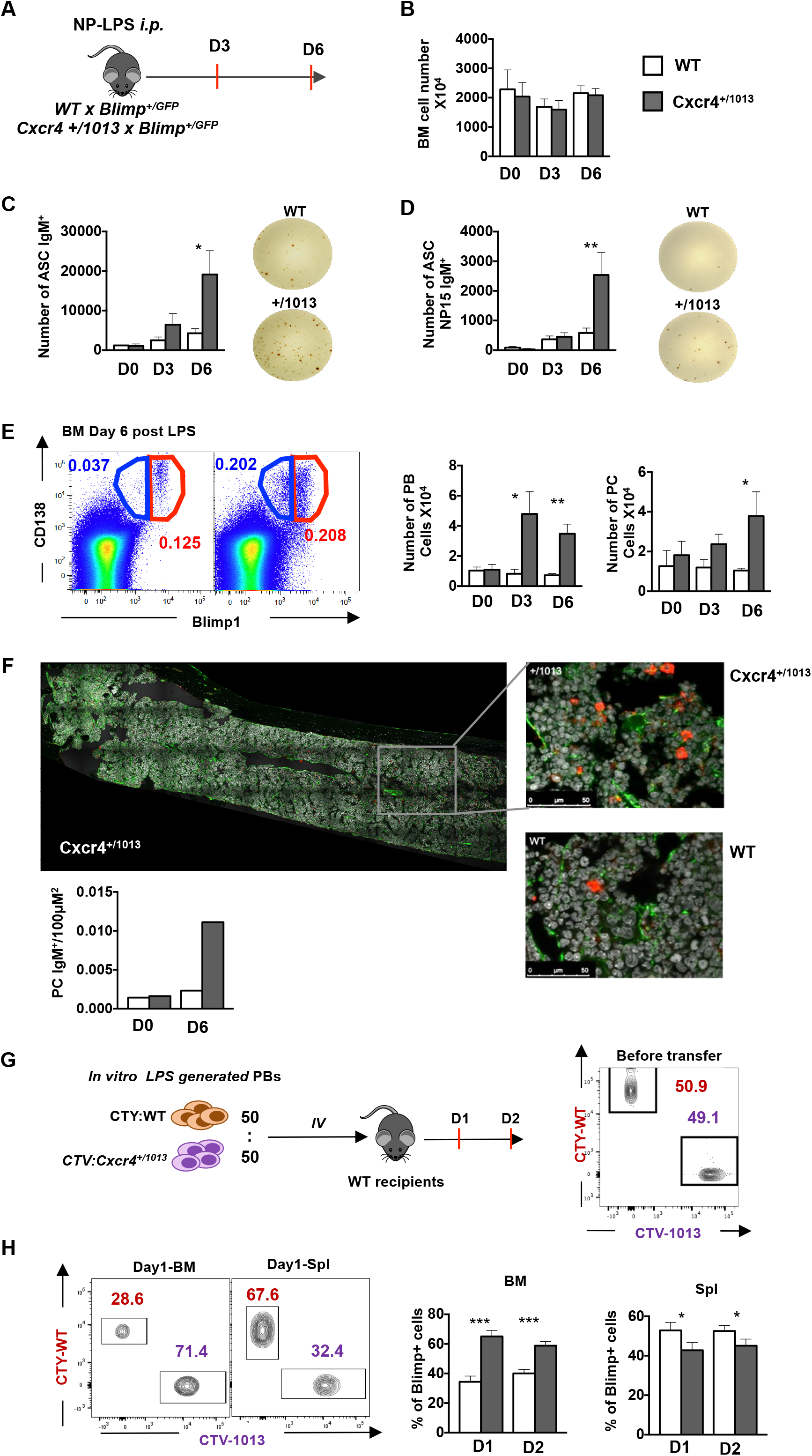
Cxcr4 desensitization limits plasmablasts accumulation within the bone marrow: **(A)** Schematic diagram describing the immunization protocol. **(B)** BM cellularity of both WT and *Cxcr4*^+/*1013*^ mice during NP-LPS immunisation. **(C-D)** Quantification and representative spots of total IgM^+^ ASCs (C) and of NP-specific IgM^+^ ASCs (D) in the BM determined by ELISpot at d0, 3 and 6 post immunization. **(E)** Representative dot plots for PC (Blimp1-GFP^hi^ CD138^+^) and PB (Blimp1-GFP^low^ CD138^+^) in the BM of both WT and *Cxcr4*^+/*1013*^ mice at day 6 post immunization (left panels). Total cell numbers for PBs and PCs at day 0, 3 and 6 are shown (right panels). **(F)** Representative images of PC staining in BM sections from WT and *Cxcr4*^+/*1013*^ mice at day 6 post immunization. Whole BM imaging is shown only for *Cxcr4*^+/*1013*^ mice. Sections were stained with Ab against laminin (green), IgM (red) and nuclei were counterstained with Hoechst 33342 (Grey) Scale bare: 40μm. The quantification of the number of IgM^+^ cells per 100μM^2^ fields is also presented. **(G)** Schematic diagram describing the transfer experiment protocol. Blimp1^GFP/+^ PBs were generated *in vitro* from splenic B cells and labelled with CTV for *Cxcr4*^+/*1013*^ cells and with CTY for WT cells. Labelled WT and mutant cells were mixed at a 1:1 ratio and transferred into WT recipients. A representative dot plot of the cell mix (1:1) prior transfer is shown (right panel). **(H)** Dot plots showing the percentages of CTV+ (*Cxcr4*^+/*1013*^ x Blimp1^GFP/+^) and CTY+ (WT x Blimp1^GFP/+^) cells in the BM and the spleen of recipient mice at day 1 post transfer (left panels). Frequency of CTV+ *(Cxcr4^+/1013^* x Blimp1^GFP/+^) and CTY+ (WT x Blimp1^GFP/+^) cells in the BM and the spleen of recipient mice at day 1 and 2 post transfer (right panel). Results are from 2-3 independent experiments (A-G) or one representative experiment out of 2 (H) (Mean ± SEM, n=2-6). Mann-Whitney U test was used to assess statistical significance (*P<0,05, ** P<0,01, ***P<0,001).

We next wondered if this phenomenon could also be observed spontaneously. We let some mice to age and observed in non-manipulated old *Cxcr4*^+/*1013*^ mice a phenotype reminiscent to the one observed in young mice after LPS injection. Old *Cxcr4*^+/*1013*^ still display a profound B cell lymphopenia mostly caused by a decreased follicular B cell compartment while the MZ compartment was preserved (Figure S4A-C). Despite this defect, a very significant increase in the frequency and number of PBs was observed in the spleen of old *Cxcr4*^+/*1013*^ mice compared to WT ones in absence of any immunization (Figure S4D-E). Moreover, the frequency and number of PCs and more particularly of PBs were also very significantly increased in the BM of mutant mice compared to the WT ones (Figure S4F-G). Altogether, these results show that in absence of Cxcr4 desensitization a chronic low-grade inflammation like the one observed with age (i.e. “inflammaging”) also promotes enhanced PB generation and aberrant BM accumulation of these cells.

Finally, we measured the intrinsic capacity of WT and mutant PBs to home to the BM by cotransfer experiments. PBs were generated *in vitro* from WT x Blimp-1 GFP and *Cxcr4*^+/*1013*^ x Blimp-1 GFP B cells stimulated for 4 days with LPS. Cells were then stained with either CTV or CTY and mixed at a 1:1 ratio (Figure 4G) before being injected *i.v*. to CD45.1 WT recipient mice. Spleen and BM homing of WT and mutant PBs were determined in each recipient mouse by flow cytometry 24 hours and 48 hours after transfer. As shown in Figure 4H, *Cxcr4*^+/*1013*^ mutant PBs had an intrinsic homing advantage to the BM compared to WT PBs at all time points considered. On the contrary, fewer mutant PBs and more WT PBs were observed in the spleen of the recipient mice (Figure 4H). This experiment suggests that mutant PBs have an intrinsic advantage in terms of BM homing. In summary, our data support that Cxcr4 desensitization is a control mechanism regulating the homing to the BM of PBs generated after LPS challenge or spontaneously in old mice experiencing inflammaging.

### Cxcr4 desensitization controls plasmablast maturation and persistence within the bone marrow

We next questioned the fate of these aberrant PBs within the BM. The analysis of a longer kinetics demonstrated that the number of PBs in the BM of *Cxcr4*^+/*1013*^ rapidly declined from day 6 and was back to the WT level 12 days after NP-LPS immunization (Figure 5A left panel). In parallel, the number of fully differentiated B220^−^ PCs progressively increased reaching a peak between 8 and 12 days after immunization and then slowly declined (Figure 5A right panel). This phenomenon was faint but detectable in the BM of WT mice while it was very pronounced in the BM of *Cxcr4*^+/*1013*^ mice. These distinct kinetics suggest that the PBs reaching the BM in absence of Cxcr4 desensitization may be able to differentiate into long-lived PCs. Moreover, we used our transfer experiments (see Figure 4H for experimental procedure) to determine whether *Cxcr4*^+/*1013*^ PBs were more likely to differentiate into long-lived PCs within the BM. At day 1, most of the transferred cells were found in the PB gate (CD138^hi^B220^+^) and very few were detected within the PC gate (CD138^hi^B220^−^) demonstrating they still had an immature phenotype (Figure 5B). At day 2 most of the transferred cells were detected amongst the CD138^hi^B220^−^ PCs while the number of PBs remained the same (Figure 5B). When we measured whether they were coming from WT (CTY+) or *Cxcr4*^+/*1013*^ (CTV+) cells we observed that around 80% of the PCs came from the *Cxcr4*^+/*1013*^ cells while the reverse was true for PBs (Figure 5C-D). Altogether, these results suggest that the Cxcr4 gain of function confers an enhanced maturation potential to BM PBs. We next assessed the fate of these cells in terms of persistence and proliferation. We observed that transferred WT and *Cxcr4*^+/*1013*^ BM PB/PC lose the cell trace dyes equivalently (Figure 5E and S5A) suggesting that they proliferate at the same rate. Moreover, we assessed BM PB and PC fate during an EdU pulse chase experiment (see Figure 3F for experimental design). At the end of the EdU labelling (6 days post immunization with NP-LPS) most of the BM PBs (CD138^+^B220^+^) were EdU^high^ suggesting that they had incorporated the dye recently (Figure 5F). In line with their proliferative nature more than 70% of BM PBs were EdU+ at day 6 at the end of the labelling (Figure 5G, top). As expected considering their quiescent nature, less than 50% of BM PCs were EdU+ after 6 days of labelling (Figure 5G, bottom). At the end of the labelling period (day 6) the frequency and number of EdU^+^ PBs and PCs were higher in *Cxcr4*^+/*1013*^ mice compared to WT ones (Figure 5F-G). This may reflect the enhanced generation of PBs in the spleen and their enhanced migration to the BM. When we followed the EdU loss during the chase we observed that both WT and mutant PBs and PCs lose it at the same rate (Figure 5G, left panels). This was also true for splenic PBs (Figure S5B). The absolute number of EdU+ PBs was more elevated in the *Cxcr4*^+/*1013*^ mice compared to the WT ones at day 6 and 8 but was not different anymore from day 12 suggesting that mutant PBs either have left the BM, died or differentiated into PCs. Interestingly, the number of EdU+ PCs was increased in the *Cxcr4*^+/*1013*^ mice compared to the WT ones at day 6, 8 and 12 (Figure 5G) hence supporting that mutant PBs may indeed give rise to PCs between day 6 and 12. Accordingly, we observed that the EdU MFI for mutant PCs was higher at day 8 than at day 6 (Figure 5F and H). As no more EdU is given at this time, this suggests that EdU^high^ PBs have given rise to EdU^high^ PCs in the *Cxcr4*^+/*1013*^ mice. This phenomenon was not observed in WT mice. Altogether, these data suggest that Cxcr4 desensitization is essential for limiting PB maturation within the BM while not affecting their proliferation status. At day 45, we did not observe any difference in terms of frequency nor absolute number of EdU+ cells in our two experimental groups. EdU+ PBs were not detected anymore while some Edu^high^ and EdU^int^ PCs were still present in the BM of both groups (Figure 5F-G). This could suggest that some PBs may have differentiated into long-lived PCs although we could not exclude that some long-lived PCs may have also incorporated EdU during the labelling. To address this point, we analysed the presence of NP-specific ASCs by ELIspot. We observed that total IgM and NP-specific ASCs were highly increased in the BM of *Cxcr4*^+/*1013*^ mice compared to the WT ones and this increase was still observed 45 days after immunization (Figure 5I). Thus, *Cxcr4*^+/*1013*^ PBs can indeed differentiate into long-lived PCs and persist for over a month in the BM. In summary, Cxcr4 desensitization limits both PB maturation into fully-differentiated PCs and PC persistence within the BM.

**Figure 5:**
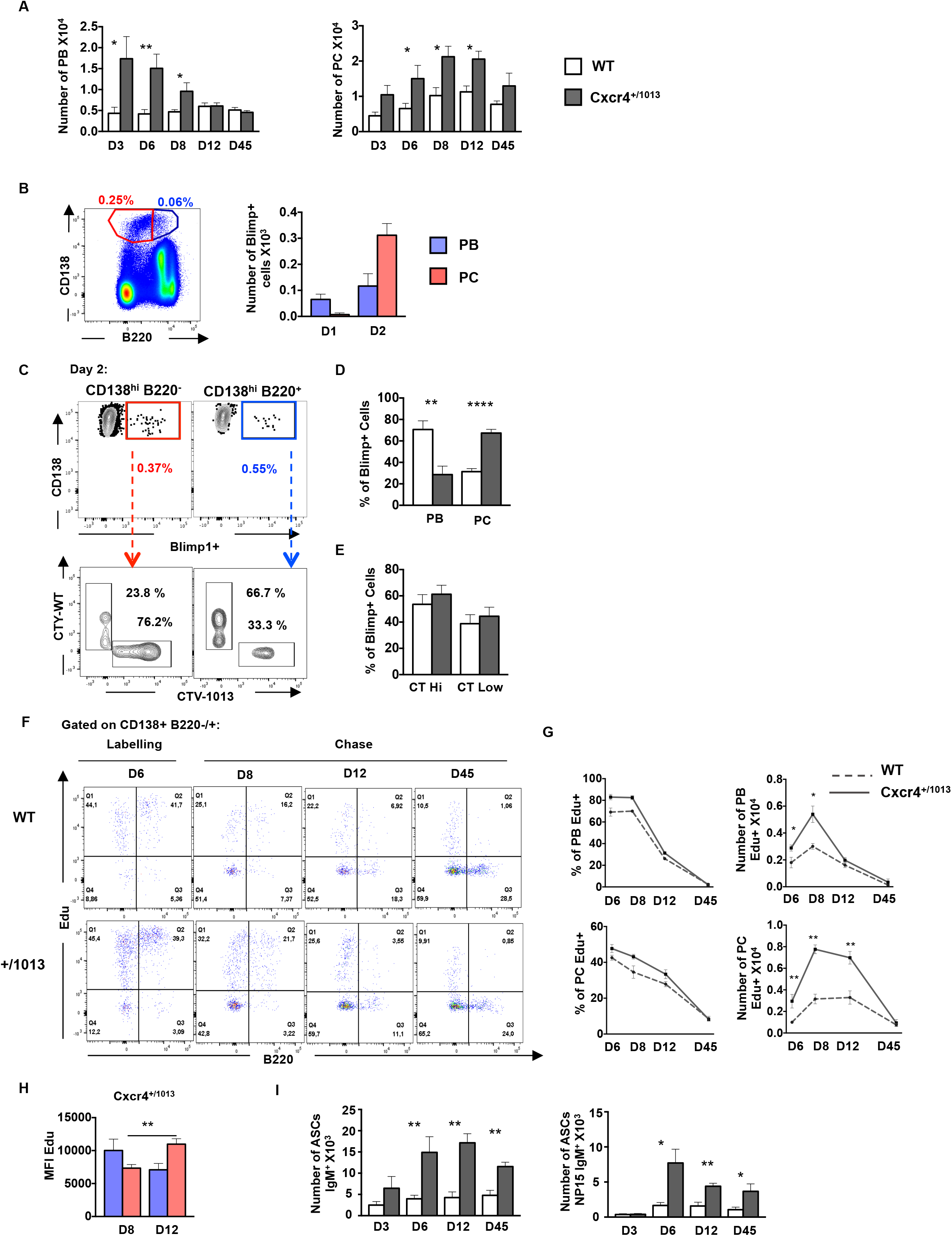
Cxcr4 desensitization controls plasmablast maturation and persistence within the bone marrow: **(A)** PCs (B220^−^CD138^+^) and PBs (B220^lo^CD138^+^) numbers in the BM of both WT or *Cxcr4*^+/*1013*^ mice post NP-LPS immunisation at the indicated time points. **(B)** Blimp1^GFP/+^ PBs were generated *in vitro* from splenic B cells and labelled with CTV for *Cxcr4*^+/*1013*^ cells and with CTY for WT cells as previously described in (Fig 4G) and transferred into WT recipient mice. Left: Representative dot plots of PCs (Red) and PBs (Blue) gating in the BM of recipient mice. Right: Total number of transferred Blimp1^GFP/+^ PBs and PCs in the BM of recipient mice at day 1 and 2 post transfer. **(C)** Left: Representative dot plots showing the frequency of CTV+ (*Cxcr4*^+/*1013*^ x Blimp1^GFP/+^) and CTY+ (WT x Blimp1^GFP/+^) cells in both the PC (CD138^+^B220^−^) and PB (CD138^+^B220^+^) compartments in recipient mice at day 2 post transfer. Right: Quantification of the frequency of CTV+ (*Cxcr4*^+/*1013*^ x Blimp1^GFP/+^) and CTY+ (WT x Blimp1^GFP/+^) in the PB and PC compartments of recipient mice at day 2 post transfer. **(D)** Heatmap showing the relative expression of PC master regulator genes presented as (2-^Δ^Ct^^) in BM PCs and PBs from both genotypes at day 6 post NP-LPS immunisation. Data are presented by applying a column Z score. **(E)** The proliferation of CTV+ (*Cxcr4*^+/*1013*^ x Blimp1^GFP/+^) and CTY+ (WT x Blimp1^GFP/+^) PB/PCs in the BM of recipient mice was assessed by measuring the dilution of the cell trace (CT) dye (no or few divisions = CT High; several divisions= CT Low) in both genotypes at day 2 post transfer. **(F)** Representative dot plots of EdU labelling of total CD138+ cells at the end of the labelling (day6) and during the chase (day 8 to 45) at the indicated time-points after NP-LPS immunisation. **(G)** Percentages and numbers of EdU+ PCs and PBs from d6 and during the chase at the indicated time-points after NP-LPS immunisation. **(H)** Analyses of EdU MFI among BM EdU+ PBs and PCs of Cxcr4^+/1013^ mice at day 8 and 12 during the chase. **(I)** Number of total ASCs IgM^+^ and NP15-IgM^+^ in the BM assessed by ELIspot at the indicated time points post NP-LPS immunisation. Results are from one representative experiment out of 2 (Mean ± SEM, n=4-5). A two-tailed Student’s t test was used to assess statistical significance (*P<0,05, ** P<0,01, ****P<0,0001).

### Cxcr4 desensitization regulates the global migratory potential of plasmablasts and plasma cells

Our data support essential roles for Cxcr4 desensitization in the control of PB maturation, of their migratory potential towards the BM and of their persistence within this organ. We thus extended our analysis by measuring the expression profile of molecules involved in cell migration and adhesion. Strikingly we demonstrated that mutant PBs and PCs from both the spleen and the BM had a unique transcriptional profile characterised by increased expression of *Cxcr4, Cxcr3, JamA, Itgal, Itgb1* and *Vcam-1* and decreased expression of *Ccr7, Cxcr5, Sell* and *Klf2* compared to WT PBs and PCs (Figure 6A-C). Actually, when we performed a principal component analysis on the basis of the expression of adhesion/ migration molecules, splenic and BM mutant PBs and PCs clustered together (Figure 6B). WT splenic and BM PBs display the more distant profile while WT BM PCs had a transcriptional profile closer to the mutant cells (Figure 6B). PC1 could be mainly accounted by the differential expression of *Cxcr5, Ccr7* and *Sell* on one hand and of *Cxcr3* and *Cxcr4* on the other hand (Figure 6C). We next measured the expression at the cell surface of selected molecules and observed that *Cxcr4*^+/*1013*^ splenic PBs expressed more Cxcr4 and less CD62L than their WT counterparts (Figure 6D). The frequency of Cxcr3+ PBs was also increased in the spleen of mutant mice compared to WT ones (Figure 6D). These differences were even more marked when we analysed PBs and PCs from the BM of WT and mutant mice (Figure 6E). In addition, mutant BM PBs express more Vla4 and Lfa1 at their surface than WT PBs. This was also observed albeit less marked for mutant and WT BM PCs (Figure 6E). Thus, in absence of Cxcr4 desensitization, splenic PBs display an enhanced homing capacity towards the BM that is not only due to the gain of function of Cxcr4 but to a more global change in the migratory code of these cells controlled by the exacerbated Cxcr4/Cxcl12 signalling. Moreover, the increased expression of several adhesion molecules observed in mutant BM PBs may explain their enhanced maturation and persistence within the BM environment.

**Figure 6:**
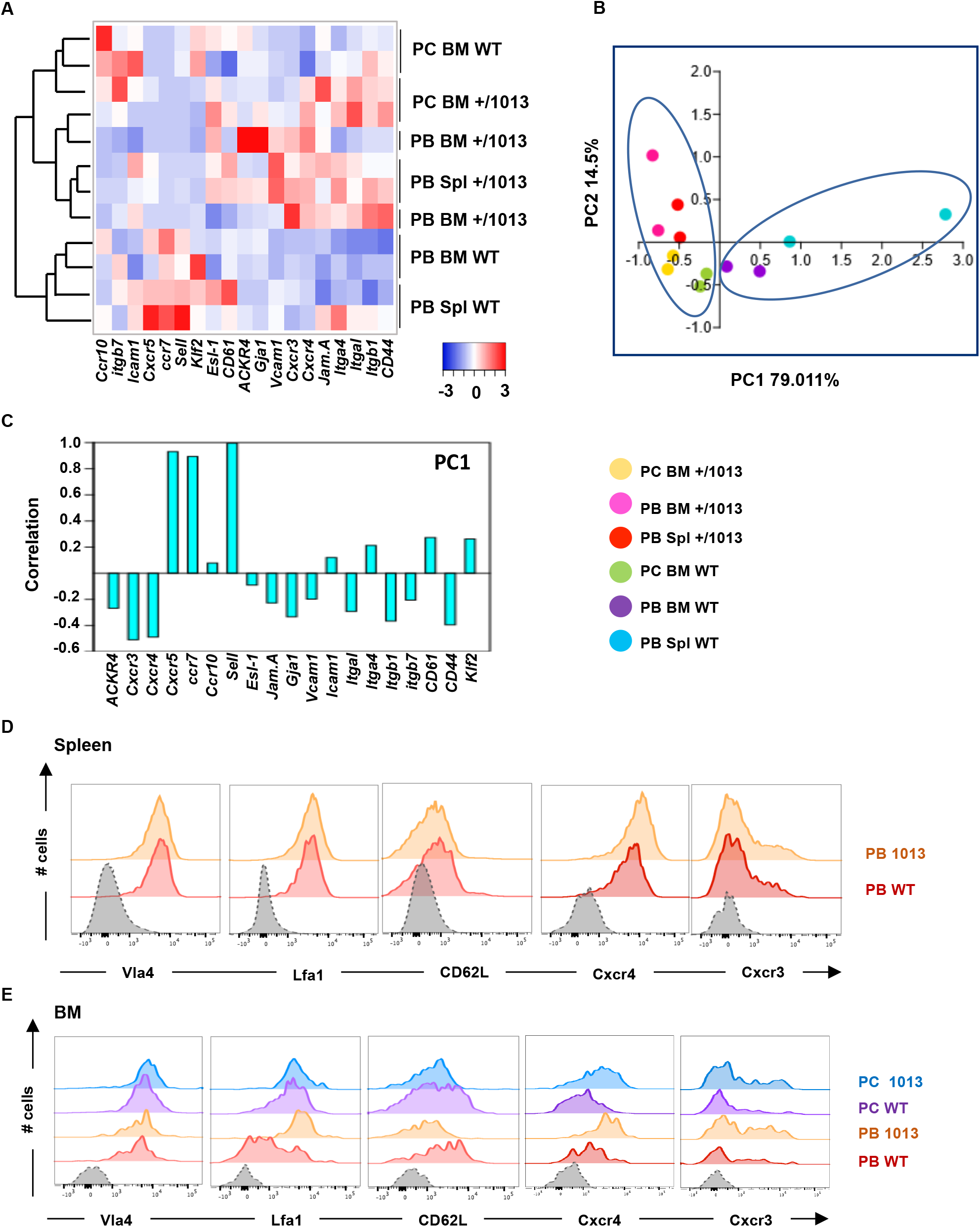
Cxcr4 desensitization regulates the migratory potential of plasmablasts and plasma cells: **(A)** Heatmap showing the relative expression of selected transcripts from BM PBs and PCs and splenic (Spl) PBs from both WT and *Cxcr4*^+/*1013*^ mice determined by Biomark multiplex qPCRs at day 6 post immunization. The heatmap was generated using the heatmapper website. Data are presented by applying a column Z score based on (2-^Δ^Ct^^). Clustering analyses was performed using a spearman rank correlation test. **(B)** Principal component analyses were performed on Spl-PBs, BM-PBs and BM-PCs from both groups using a variance-covariance test. **(C)** Gene variance correlations in the principal component analysis 1 (PCA1) are shown. **(D)** Representative histograms for the indicated proteins from Spl PBs from both WT and *Cxcr4*^+/*1013*^ mice are shown. **(E)** Representative histograms for the indicated proteins from BM PBs and PCs from both WT and *Cxcr4*^+/*1013*^ mice are shown. Results are from one representative experiment out of 2 (n=2 for A-C; n=3-4 for D-E).

## Discussion

Cxcr4 is a well-known regulator of the immune system. However, how the fine-tuning of its signalling may impact on homeostatic processes is still poorly understood. Here we demonstrated that the desensitization of Cxcr4 is an essential regulatory mechanism controlling the B cell immune response upon TLR-mediated activation. In particular, we demonstrated a crucial role for Cxcr4 desensitization in limiting follicular B cell entry into cycle. This finding could have important implications for B cell homeostasis but also for the development of B cell lymphomas. On the homeostatic side, this suggests that exacerbated Cxcl12/Cxcr4 signalling may promote B cell cycling in other contexts. One can think of the GC dark zone in which Cxcl12 is highly expressed by specific stromal cells. This enhanced production of Cxcl12 might contribute to the high cycling of B cells observed in this anatomical site. Moreover, previous studies have linked B cell division to their capacity to differentiate into PCs and thus the enhanced cycling of follicular B cells may contribute to their enhanced differentiation both *in vivo* and *in vitro*. At the pathological level, Cxcl12 is highly expressed in several lymphomas^21^ including WM^22^ and this may contribute to an accelerated G0/G1 transition. Interestingly, an enhanced CXCR4 expression was reported in most WM patients, even those not bearing the *CXCR4* gain of function mutation^23,24^. Thus, even in the non-mutated patients, the CXCR4/CXCL12 signalling axis may contribute to the pathology.

Another observation we made was that Cxcr4 desensitization limits the differentiation of MZ B cells into PBs independently of cell cycle. Although the molecular mechanism at play is still not clear, fine-tuning of Cxcl12/Cxcr4 signalling appears to be necessary for controlling the balance between the expression of the “pro-PC” transcription factor Blimp1 and of the “anti-PC” transcription factor Bach2 in MZ B cells. Further work will be required to unravel how Cxcl12/Cxcr4 signalling impacts on this balance and how this translates into enhanced PB differentiation.

Our data also suggest that one of the main functions of Cxcr4 desensitization in B cell immune response is to control PC homeostasis. PB differentiation is enhanced upon Myd88 activation in absence of Cxcr4 desensitization but the implication of this process goes beyond limiting their generation. Indeed, PBs and PCs bearing the *Cxcr4* gain of function mutation have distinct migratory and maturation potentials than their WT counterparts. Interestingly, our results suggest that this enhanced BM tropism is not only due to the gain of function of Cxcr4 *per se* as the expression of a whole range of migration/adhesion factors are modified in PBs and PCs in absence of Cxcr4 desensitization. Moreover, Cxcr4 desensitization also controls PB maturation into fully differentiated PCs. Whether this is due to a direct impact of the exacerbated Cxcl12/Cxcr4 signalling on the maturation transcriptional program or whether it is an indirect effect induced by different microenvironmental cues specific to mutant PB/PCs remains to be demonstrated.

Conceptually our results may suggest an important role for Cxcr4 desensitization in limiting the laps of time during which Abs generated in the course an extra-follicular response are produced. Indeed, Cxcr4 desensitization not only dampens the early generation of PBs in the secondary lymphoid organs after immunization but also limits their homing to and maturation within the BM, hence ensuring that only few long-lived PCs generated during the extrafollicular response persist. As such, Cxcr4 desensitization may be a gatekeeper ensuring that the Ab produced by PBs during the extrafollicular response only form a transient wave of protection. Furthermore, limiting PB homing to the BM may also limit the filling of BM niches normally dedicated to other cell types, including GC-derived high affinity PCs that start arising from day 9 after immunization^25^.

Another surprising result was the small but consistent effect of Cxcl12 on its own on PC differentiation and B cell cycling in absence of Cxcr4 desensitization. Although this effect was not of a magnitude comparable to the differentiation and proliferation induced by TLR ligands it was always observed and was totally abolished by a Cxcr4 antagonist hence confirming that it was indeed caused by the exacerbated Cxcl12/Cxcr4 signalling. Endotoxin-low media were used in all *in vitro* experiments suggesting this is not the result of a contamination with small doses of LPS. This observation suggests that unless controlled by the desensitization of the receptor, this axis may act as a mitogen. It is thus somehow surprising that the addition of Cxcl12 did not significantly enhance the CpG- and LPS-mediated differentiation and cycling observed for *Cxcr4*^+/*1013*^ B cells. Several mechanisms could explain this observation. First, we cannot absolutely exclude that some Cxcl12 was produced by the cultured splenocytes independently of any exogenous addition and that these suboptimal doses were sufficient to explain the differences observed between WT and *Cxcr4*^+/*1013*^ samples. However, we did not detect any expression of *Cxcl12* by qPCR in splenic B cells. Furthermore, we analysed Cxcl12-dsRed reporter mice and could not detect any dsRed+ splenic B cells strongly supporting that these cells cannot produce Cxl12 themselves. Previous report suggested that CXCR4 could be part of the LPS “sensing-apparatus” together with TLR4 and CD14^26,27,26,28^. However, this is not the case for CpG stimulation for which we also observed an enhanced B cell differentiation in absence of Cxcr4 desensitization. Alternatively, it is also possible that the Cxcr4 gain of function imprints B cells with specific abilities *in situ* following binding of its ligand that they conserve *ex vivo*. In addition, we previously reported that Cxcl12 expression and detection was increased in the spleen of *Cxcr4*^+/*1013*^ mice compared to controls^29^. Recent work in our laboratory also reports that Cxcl12 is increased in the BM of mutant mice (Nguyen *et al*. under preparation). In line with that hypothesis, recent studies on human and mouse B cells revealed that PC differentiation is coupled to division-dependent DNA demethylation and PC gene regulation^30–32^. Together with the results presented here, these observations suggest that the enhanced Cxcl12/Cxcr4 signaling mediated by the gain of function of the receptor together with the enhanced availability of its ligand, may contribute to a deep imprinting of the B cells that could partly explain the phenotypes observed. Further work will be required to establish whether the strength of the Cxcr4-induced signalling promotes a specific transcriptional signature or epigenetic imprinting on the different B cell subsets.

TLR4 and TLR9 signalling that promoted mutant B cell differentiation in our study via LPS and CpG stimulation respectively, both rely on the adaptor MyD88. This is of particular interest considering that almost all WM patients present a *MYD88* mutation and 20 to 40% also present a *CXCR4* mutation^15,33^. Previous work on WM cell lines expressing the mutated forms of both CXCR4 and MYD88 suggest that the signals induced by both pathways add-up rather than synergize by inducing differently several kinases in particular BTK and AKT^34^. As a consequence, the presence of both mutations is associated with increased resistance to several kinase inhibitors including Ibrutinib, recently approved by the FDA for the treatment of WM^34,35^. Moreover, *CXCR4* mutated patients present increased BM involvement and serum IgM leading to enhanced hyperviscosity and symptomatic disease^14^. Our results show that the exacerbated Cxcl12/Cxcr4 signalling occurring in absence of Cxcr4 desensitization promotes the homing of immature PBs to the BM and once there favours their maturation and persistence. It is thus possible that a similar mechanism takes place in WM and contributes to the BM homing of malignant cells and to the niche-mediated therapy resistance previously reported^36^.

## Material and Methods

### Mice

*Cxcr4*^+/*1013*^ mice (C57Bl/6J background) were generated as previously described^13^. Blimp^GFP/+^ (C57Bl/6J background) mice were obtained from Dr. Stephen Nutt (The Walter and Eliza Hall Institute of Medical Research, Australia)^37^ and crossed with *Cxcr4*^+/*1013*^ mice. Mice between 8-16-week-old or 1-year old (old cohort, Figure S4) were used. Mice were bred into our animal facility under 12-h light/dark cycles, specific-pathogen-free conditions and fed *ad libitum*. All experiments were conducted in compliance with the European Union guide for the care and use of laboratory animals and has been reviewed and approved by an appropriate institutional review committee (C2EA-26, Animal Care and Use Committee, Villejuif, France).

### Immunization and Edu feeding

Mice were immunized intra-peritoneally with 25 μg of 4-hydroxy-3-nitrophenylaceyl-lipopolysaccharide (NP-LPS) (Biosearch Technologies). Blood, Spleen and BM were harvested and analysed 3 and 6 days after challenge. For EdU pulse-chase assays, mice were immunised with 25 μg NP-LPS and injected intra-peritoneally with 1 mg Edu (Sigma-Aldrich) and maintained on drinking water containing 500 μg/ml Edu and 1% glucose during 6 days. The Edu labelling was followed by a chase period of 2, 6, 12 and 33 days (6, 8, 12 and 45 days post NP-LPS immunisation). PCs and B cells were detected by flow cytometry by combining surface staining with intracellular staining using the Edu-labelling kit following the manufacturer’s instructions (Sigma, EdU-Click 488/Base click).

### Single cell suspension

Isolation of organs was performed upon euthanasia by inhalation of CO2. The BM was harvested from femurs, tibias and hip by centrifugation in culture medium (RPMI with 10% decomplemented foetal calf serum and penicillin 100Units/mL/streptomycin100μg/mL (all from Gibco)). Spleens were gently mashed on a 70μm nylon mesh and recovered cells were then suspended in culture medium. Red blood cells from spleen and BM single cell suspension were lysed for 5 min in 1 mL ammonium-chloride-potassium buffer. Harvested cells from tissues were either directly immunophenotyped or used for *in vitro* functional assays.

### Adoptive transfer experiments

Splenocytes from both WT x Blimp1^GFP/+^ and *Cxcr4*^+/*1013*^ x Blimp1^GFP/+^ mice were stimulated *in vitro* (see section *in vitro* assays) with the indicated LPS concentration for 4 days. Cells from each genotype were loaded with either CTV (Cell trace violet) or CTY (Cell trace yellow) according to manufacturer’s instructions (Thermofisher). A 50:50 mix of Blimp1^GFP/+^ cells of both genotypes was made and checked by flow cytometry prior injection. 10^6^ cells were transferred intravenously (i.v.) in C57Bl/6J CD45.1 WT recipients. Blood, BM and spleens were harvested and analysed at 4, 24 and 48 hours post transfer.

### *In vitro* functional assays

For PB differentiation, B cells were isolated from total splenocytes using naïve mouse B-cell isolation kit (Miltenyi Biotec) according to the manufacturer’s instructions. Total splenocytes or isolated B cells were cultured using RPMI (with 10% decomplemented foetal calf serum, 1% penicillin 100Units/mL/streptomycin100μg/mL (Gibco), 50μM β-mercaptoethanol (PAN biotech), 1mM sodium pyruvate (Gibco) and Minimum essential medium non-essential amino acids (MEM NEAA) 1X (Gibco)) and stimulated with 1 U/mL of IL-4 (Miltenyi) and 5 ng/mL IL-5 (Miltenyi) supplemented with 1μg/mL LPS (Invivogen) or 10μg/mL CpG ODN (Invivogen) and/or 50 nM Cxcl12 (R&D) and/or 10μM AMD3100 (Sigma Aldrich) for up to 5 days at 37°C. PB differentiation was then assessed by flow cytometry with the Abs listed in table (1). For apoptosis assays, apoptosis was measured with the PE Annexin-V Apoptosis Detection Kit I (BD Pharmingen) according to manufacturer’s instructions or intra cellular staining was performed for V450-anti-cleaved caspase 3 (BD). For proliferation, total splenocytes were loaded with Cell trace violet (CTV) (thermofisher) according to manufacturer’s instructions and then stimulated as previously described. CTV dilution was assessed by flow cytometry. For cell cycle analysis, cells from different stimulation conditions were permeabilized and fixed according to the manufacturer’s instructions with the FOXP3 permeabilization kit (Foxp3/Transcription Factor Staining Buffer Set ebioscience) and then labelled with an anti-Ki-67 antibody (clone B56, mouse IgG1; BD) and DAPI was added before flow cytometry analysis.

### Flow cytometry

Single cell suspensions were stained as previously described (biajoux et al 2016), using the Abs described in Supplementary Table 1. Analyses were carried out on an LSRII Fortessa flow cytometer (BD Biosciences) and data were analyzed with the Flowjo software (TreeStar, Ashland, OR). NP conjugated to PE was from Biosearch Technologies. Live/Dead Fixable Yellow, Aqua or red Dead Cell Stain Kits (Biolegend) were used.

### Immunofluorescence

Spleens were fixed for 4 hours in PBS-paraformaldehyde (PFA) 4% and were subsequently washed in PBS and left over night in PBS with 20% sucrose. Spleens were then embedded in OCT (TissueTek) and stocked at −80°C before sectioning. Cryosections (7 μm) were permeabilized with PBS triton X100 0.5% for 15 min, washed twice with PBS and blocked with PBS BSA 5% for 1 hour. Sections were stained with primary Abs (rabbit anti-mouse laminin (Sigma Aldrich), Alexa fluor 594 Goat anti-mouse IgM (Thermofisher)) for 1 hour at room temperature (RT). Sections were washed twice with PBS and incubated with the appropriate secondary Ab (Alexa fluor 488 goat anti-Rabbit (Thermofisher)) and Hoechst 33342 for 1 h at RT. Mounting was done using Vectashield medium (Vector Laboratories). Slides were scanned using a NanoZoomer Digital Pathology system using X40 objective lense with numerical aperture 0.75 (Hamamatsu Photonic). Bone marrow embedding, sectioning and staining were performed as described^38^. Briefly, freshly dissected bones were fixed in PBS-PFA 4% overnight followed by one-week decalcification in EDTA (0.5M) at pH 7.4. Bones were then incubated in PBS 20% sucrose and 2% polyvinylpyrrolidone (PVP) at 4°C overnight. The embedding was performed using PBS with 20% sucrose, 2% PVP and 8% gelatin. Bones were sectioned (30 μm) using a cryostat. Sections were rehydrated in PBS 1X, incubated 20 min at RT in PBS with 0.3% triton X-100, saturated in blocking solution (PBS-BSA 5%) for 1 hour and finally incubated with primary Abs: rabbit anti-mouse laminin (Sigma), Alexa fluor 594 Goat anti-mouse IgM (Thermofisher) for 1 hour at RT. After washing, sections were incubated with the appropriate secondary Abs: Alexa fluor 488 goat anti-Rabbit (Thermofisher) for 1h at RT with Hoechst 33342 for nuclear staining. Mounting was done using Permafluor mounting medium (Thermofisher). Images were acquired using TCS SP8 confocal microscope and processed with Fiji.

### ELISA and ELISpot

Sera were harvested by tail bleed or by cardiac puncture after euthanasia and IgM or anti-NP IgM titers were determined by ELISA. Sera were incubated 2 h at 37°C on wells pre-coated with 5 μg/mL of NP15-BSA or goat-anti-mouse IgM (Southern Biotech) and saturated with PBS 2% BSA. The plates were then washed and incubated with peroxidase-conjugated goat anti-mouse IgM Fc-specific (Southern Biotech). In differentiation assays, culture supernatants were incubated 2 h at 37°C on wells pre-coated with 5 μg/mL of goat-anti-mouse IgM (Southern Biotech) and saturated with PBS 2% BSA. Plates were then washed and incubated with peroxidase-conjugated goat anti-mouse IgM (Southern Biotech). Detection was done with TMB Substrate Reagent Set (BD OptEIA) according to manufacturer’s instructions. The plates were read at OD405 nm with a Mithras LB 940 Multimode Microplate Reader (Berthold Technologies). Enumeration of total IgM and NP-specific IgM ASCs were performed by incubating 1×10^5^ or 5×10^4^ spleen or BM cells per well overnight at 37°C on plates precoated with NP15-BSA or goat anti-mouse IgM Fc-specific at 5 μg/mL and saturated with culture medium containing 10% FCS. Plates were then washed and incubated with peroxidase-conjugated goat anti-mouse IgM (Southern Biotech). Spots were revealed with AEC (Sigma-Aldrich). The reaction was stopped with water and analyzed using an AID ispot Reader (Autoimmun Diagnostika GMBH).

### Time lapse imaging

B cells were isolated as previously described (see section *in vitro* assays) and stimulated with LPS, IL4 and IL5 at the indicated concentrations. Cells were either imaged using a Biostation NIKON in a 37°C, 5% Co2 supplemented chamber at a concentration of 10^4^/mL or using the Incucyte^®^ S3 at a concentration of 10^4^ cell per well. Cells were imaged for phase-contrast and GFP fluorescence each 15 or 30 minutes. Images and movies where analysed with the Incucyte S3 program, the biostation IM programme and Fiji.

### Multiplex qPCR

Multiplex qPCR analysis was performed using the Biomark system. PCs and PBs (B220^−^ CD138^+^ and B220^lo^CD138^+^ respectively) or FoB, MZB or total B cells (B220^+^CD19^+^ CD23^+^ CD21^lo^, B220^+^CD19^+^CD23^−^CD21^+^ and B220^+^CD19^+^ respectively) were sorted at 100 cells/wells directly into PCR tubes containing 5μl of reverse transcription/pre-amplification mix, as previously described^39^. Briefly, the mix contained 2X Reaction mix and superscriptIII (CellDirect One-Step qRT–PCR kit, Invitrogen) and 0,2X Taqman assay (Life technologies) (Supplementary Table 2). Targeted cDNA pre-amplification was performed during 20 cycles and the pre-amplified product was diluted 1:5 in TE buffer before processing with Dynamic Array protocol according to the manufacturer’s instructions (Fluidigm). Cells expressing *Gapdh, Actb* and control genes (*Prdm1, Irf4* and *Xpb1* for plasma cells and *Pax5* for B cells) and not negative control (*CD3e*) were considered for further analysis. Mean expression of *Actb* and *Gapdh* gene was used for normalization. Heat maps were generated with (http://www.heatmapper.ca) using the Z scores and PCA with Past 3.X software using the variance-covariance test.

### qRT-PCR

Total cellular RNA was extracted from samples using the RNeasy Plus Mini kit (Qiagen) and was quantified using NanoDrop technology (Thermo Scientific, Wilmington, DE). RNA was reverse transcribed with pd(T)-15 (Roche) and Moloney Murine Leukemia Virus reverse transcriptase (Invitrogen). Amplification of cDNAs was performed by quantitative real-time PCR reactions on a Light Cycler instrument (LC480, Roche Diagnostics) with the Light Cycler 480 SYBR Green detection kit (Roche Diagnostics) using the primers listed in Supplementary Table 2. β-actin (*Actb*) was used as the reference standard for normalization and relative quantification of fold differences in mRNA expression was determined by the comparative delta-delta-CT (2^−_ΔΔ^CT^_^) method.

### Statistical analysis

Data are expressed as means ± SEM. The statistical significance between groups was evaluated using the two-tailed Student’s t test or the two-tailed unpaired Mann-Whitney non-parametric test (Prism software, GraphPad). Heatmaps were presented using the Z score and clustering analyses were performed using the spearman rank correlation test (The heatmapper http://www.heatmapper.ca).

## Author contribution

NA performed experiments, analysed data and wrote the manuscript. AB performed experiments and analysed data. JN and VR helped with experiments. KB contributed to the project design and to the manuscript redaction. ME designed the project, analysed data and wrote the manuscript.

## Acknowledgements

We thank M-L. Aknin, Dr. H. Gary, F. Gaudin, V. Nicolas, B. Lecomte (IPSIT, technical facilities PLAIMMO, PHIC, MIPSIT and AnimEx, Clamart) for their technical assistance. The study was supported by the Laboratory of Excellence in Research on Medication and Innovative Therapeutics (LabEx LERMIT) (ME and KB), an ANR @RAction grant (ANR-14-ACHN-0008) and a Fondation Arthritis grant to ME, and an ANR PRC (ANR-17-CE14-0019) for KB. NA, AB, VR, JN, KB and ME are members of the LabEx LERMIT supported by ANR grant ANR-10-LABX-33 under the program “Investissements d’Avenir” ANR-11-IDEX-0003-01. NA was supported by a PhD fellowship from the French Ministry for education and by a 4^th^ year PhD fellowship from the Fondation ARC. VR was supported by a PhD fellowship from the FRM. JN was supported by a PhD fellowship from the DIM cancéropole and by a 4^th^ year PhD fellowship from the FRM.

---

**Supplementary figure 1:**
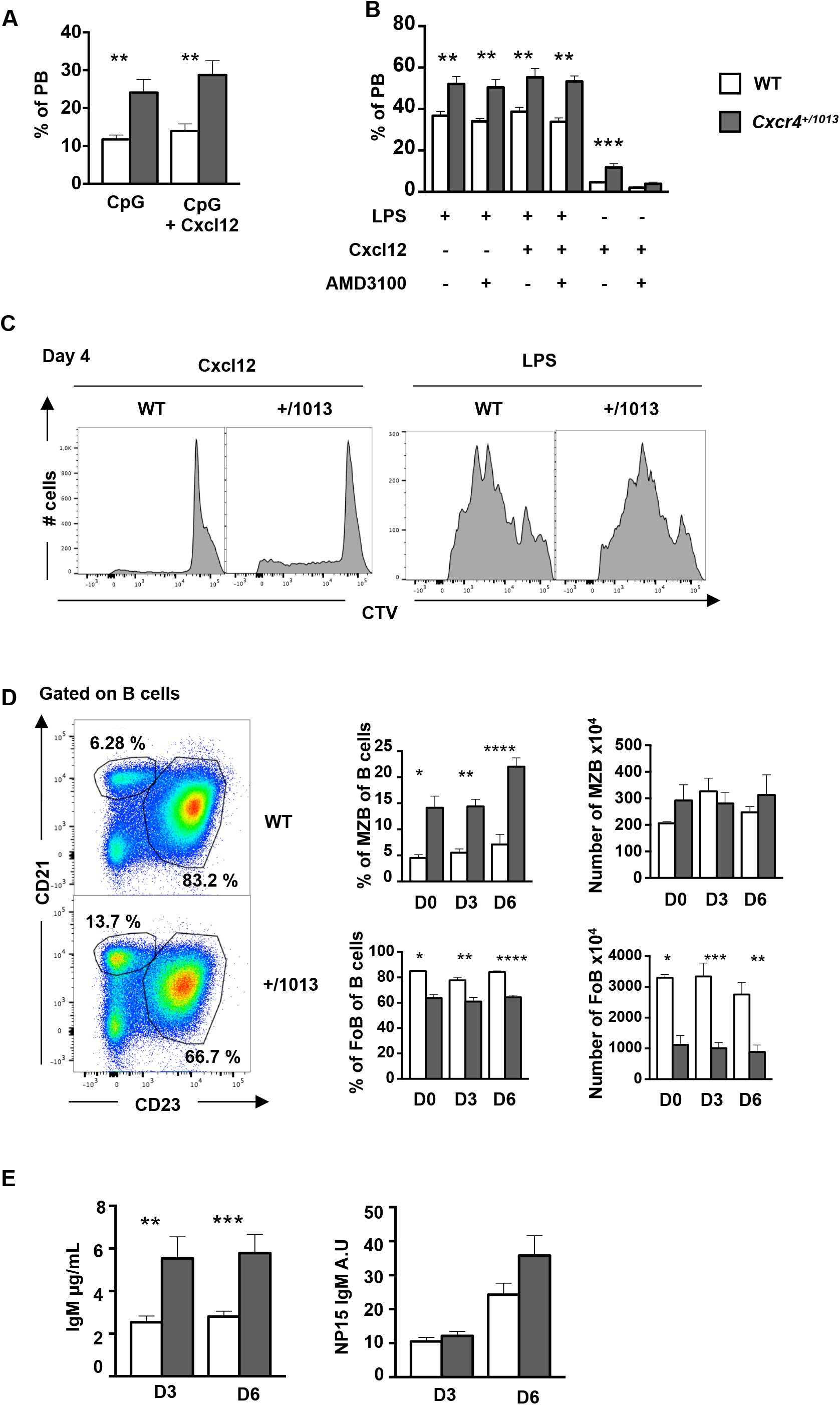
The gain of function of Cxcr4 impacts the B cell compartment and enhances MyD88-dependent PC differentiation: **(A)** Splenic B cells were cultured in presence of CpG or CpG+Cxcl12 for 4 days, the percentage of PBs was assessed by FACS **(B)** Splenocytes were cultured in presence of LPS and/or Cxcl12 and/or AMD3100. The frequency of PBs generated was assessed by FACS. **(C)** Representative histograms showing CTV dilution after 4 days of either Cxcl12 or LPS stimulation. **(D)** Left panel: representative dot plots of the gating strategy for MZB (CD21^+^CD23^−^) versus FoB (CD21^lo^CD23^+^) cells among total B cells. Right panel: percentage and total number of both MZB and FoB cells from both WT and *Cxcr4*^+/*1013*^ mice at day 3 and 6 post LPS immunization. **(E)** Serum titers of both total IgM^+^ and NP15-IgM^+^ from both genotypes were measured by ELISA at day 3 and 6 post immunization. Results are from 2-4 independent experiments (D-E) (Mean ± SEM, n=3-4). Mann–Whitney U test (A-B-E) or two-tailed Student’s t tests (D) were used to assess statistical significance (*P<0,05, ** P<0,01, *** P<0,001, ****P<0,0001).

**Supplementary figure 2:**
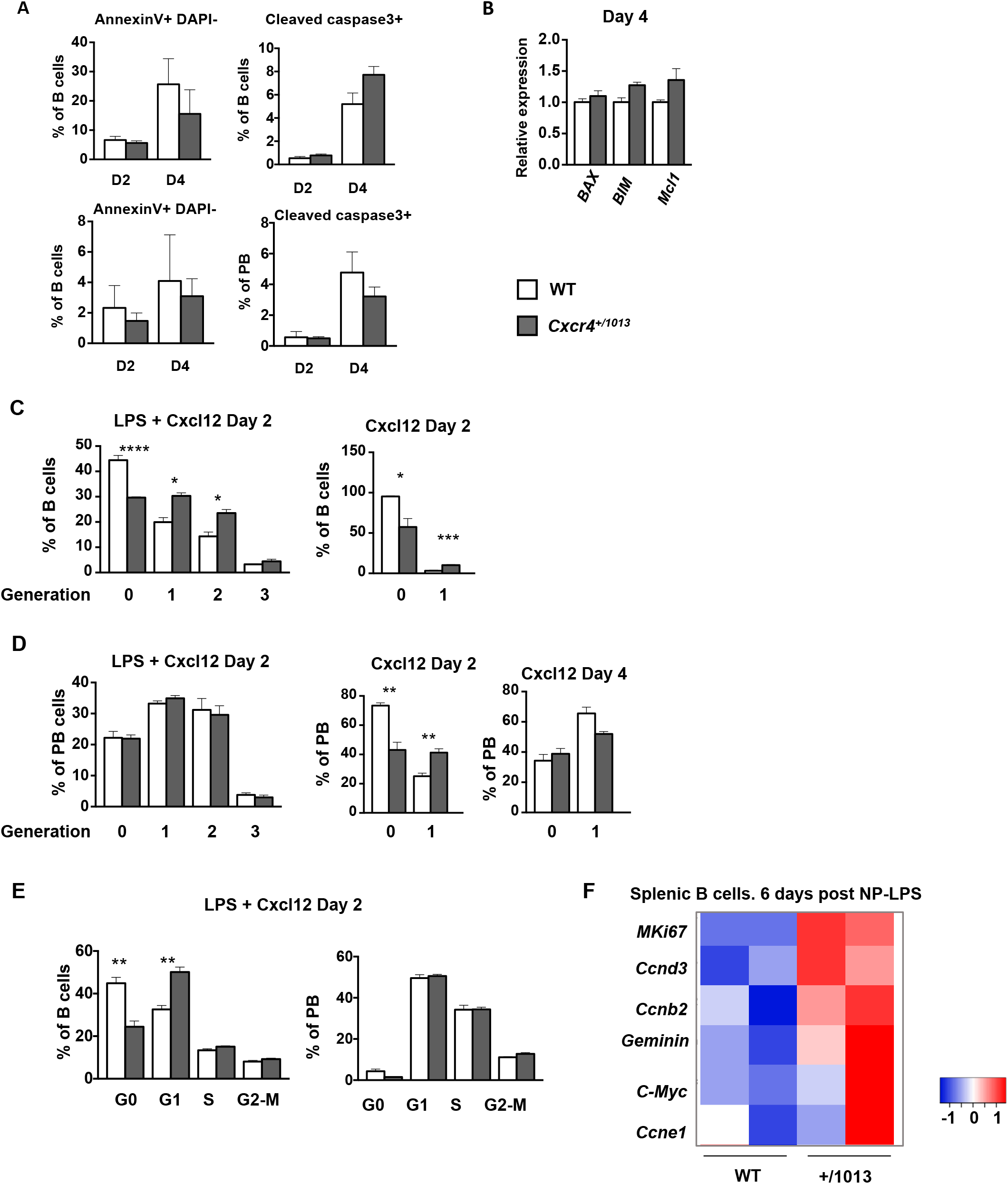
Cxcr4 desensitization controls LPS-mediated B cell cycling but not apoptosis: **(A-B)** Splenocytes from WT and *Cxcr4*^+/*1013*^ mice were cultured in presence of LPS for 4 days. **(A)** Apoptosis was assessed by measuring the frequency of Annexin V and cleaved caspase 3 positive B cells (top) and PBs (bottom) from both genotypes at the indicated time points. **(B)** Expression of *BAX, BIM* and *Mcl1* was measured by qPCR on cDNA from cells from both genotypes cultured for 4 days. Expression levels were normalized to the level of *Actb* transcripts. The fold change compared to WT B cell expression is shown. **(C-D)** Splenocytes from WT and *Cxcr4*^+/*1013*^ mice were loaded with CTV and cultured in presence of LPS +/− Cxcl12 for 2 days. The frequency of B cells and PBs present in each generation based on the CTV-dilution at day 2 and/or 4 is shown. **(E)** Frequency of splenic B cells and PBs in each cell cycle phase at day 2 post LPS+Cxcl12 stimulation. **(F)** Heatmap showing the relative expression of different mRNA from WT and *Cxcr4*^+/*1013*^ splenic B cells determined by Biomark multiplex qPCRs at day 6 post immunization. Results are from one representative experiment out of 2 (Mean ± SEM, n=2-3). A two-tailed Student’s t tests was used to assess statistical significance (*P<0,05, ** P<0,01, ****P<0,0001).

**Supplementary figure 3:**
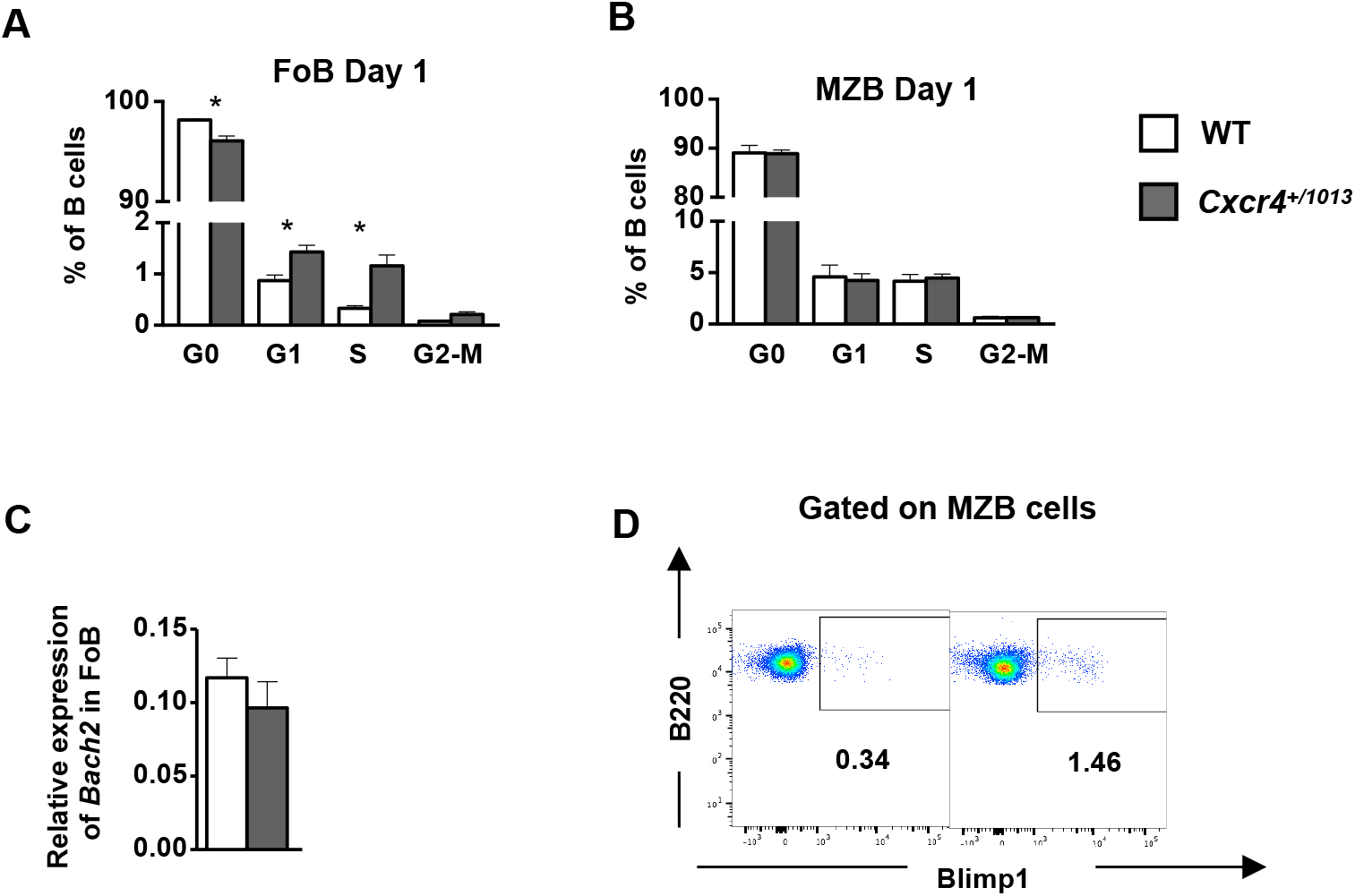
Cxcr4 desensitization differentially controls cell cycling and differentiation in follicular and marginal zone B cells: **(A)** Frequency of FoB cells present in each cell cycle phases at day 1 post LPS stimulation. **(B)** Frequency of MZB cells present in each cell cycle phases at day 1 post LPS stimulation. (C) Relative expression of *Bach2* in FoB cells presented as (2-^Δ^Ct^^). **(D)** Representative dot plots for Blimp1/GFP expression in MZB cells in the spleen of both WT and *Cxcr4*^+/*1013*^ mice at day 6 post immunization. Results are from one representative experiment out of 2 (Mean ± SEM, n=3). A two-tailed Student’s t tests was used to assess statistical significance (*P<0,05).

**Supplementary figure 4:**
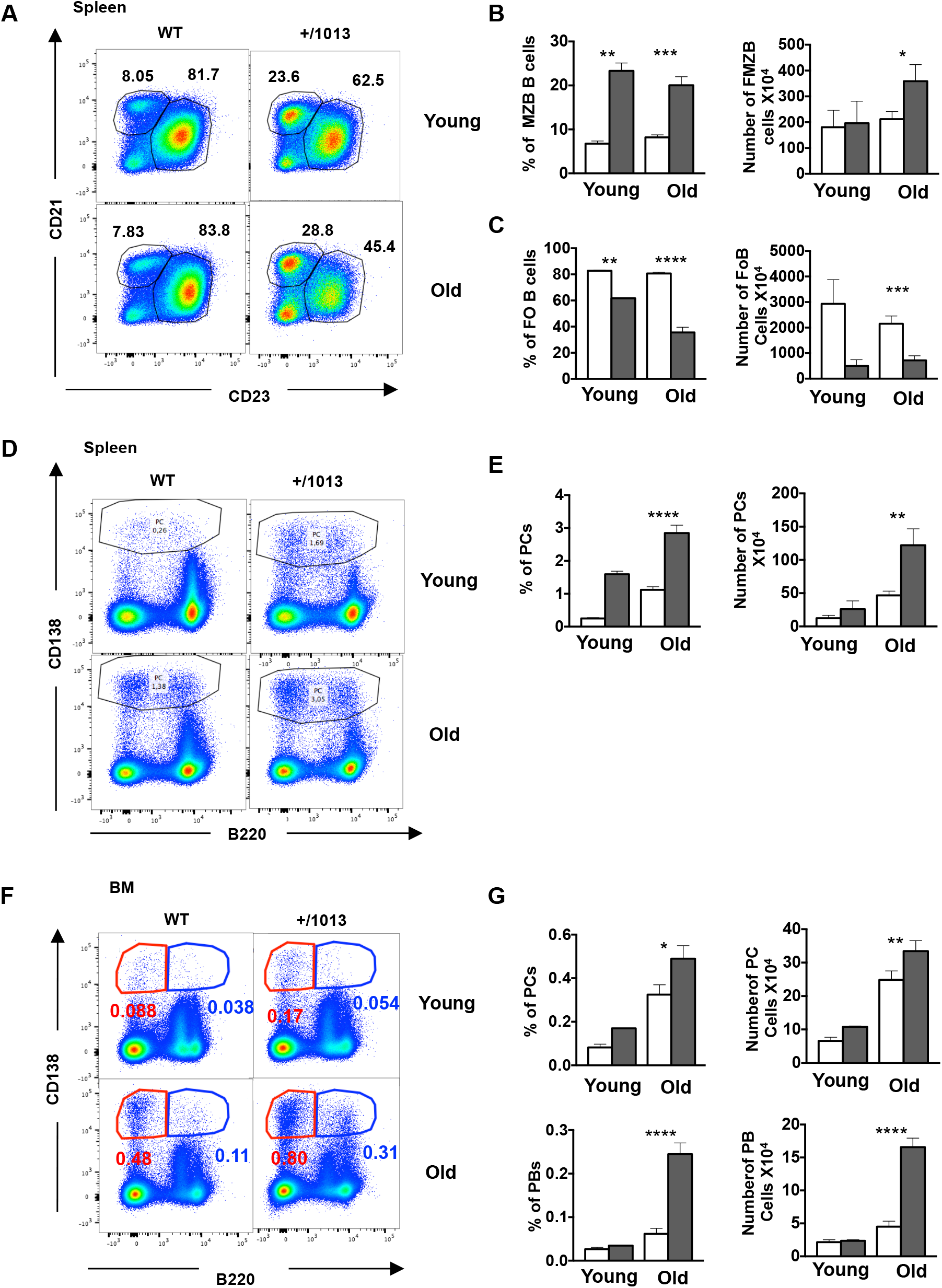
Cxcr4 desensitization blocks the spontaneous generation and bone marrow homing of plamablasts and plasma cells in old mice: **(A)** Representative dot plots showing the gating strategy for MZB and FoB cells in the spleen of young and old WT and *Cxcr4*^+/*1013*^ mice. **(B)** Frequency and total number of MZB cells in young and old mice from both genotypes. **(C)** Frequency and total number of FoB cells in young and old mice from both genotypes. **(D)** Representative dot plots showing the gating strategy for PCs in the spleen of old and young mice of both genotypes. **(E)** Frequency and total number of PCs in the spleen of young and old mice. **(F)** Representative dot plots for PBs (B220^lo^ CD138^+^) and PCs (B220^−^CD138^+^) in the BM of young and 1-year-old mice from both genotypes. **(G)** Frequency and total number of PBs and PCs in the BM of young and old mice from both genotypes. Results are from 2 independent experiments (Mean ± SEM, n=3-5). A two-tailed Student’s t tests was used to assess statistical significance (*P<0,05, ** P<0,01, *** P<0,001, ****P<0,0001).

**Supplementary figure 5:**
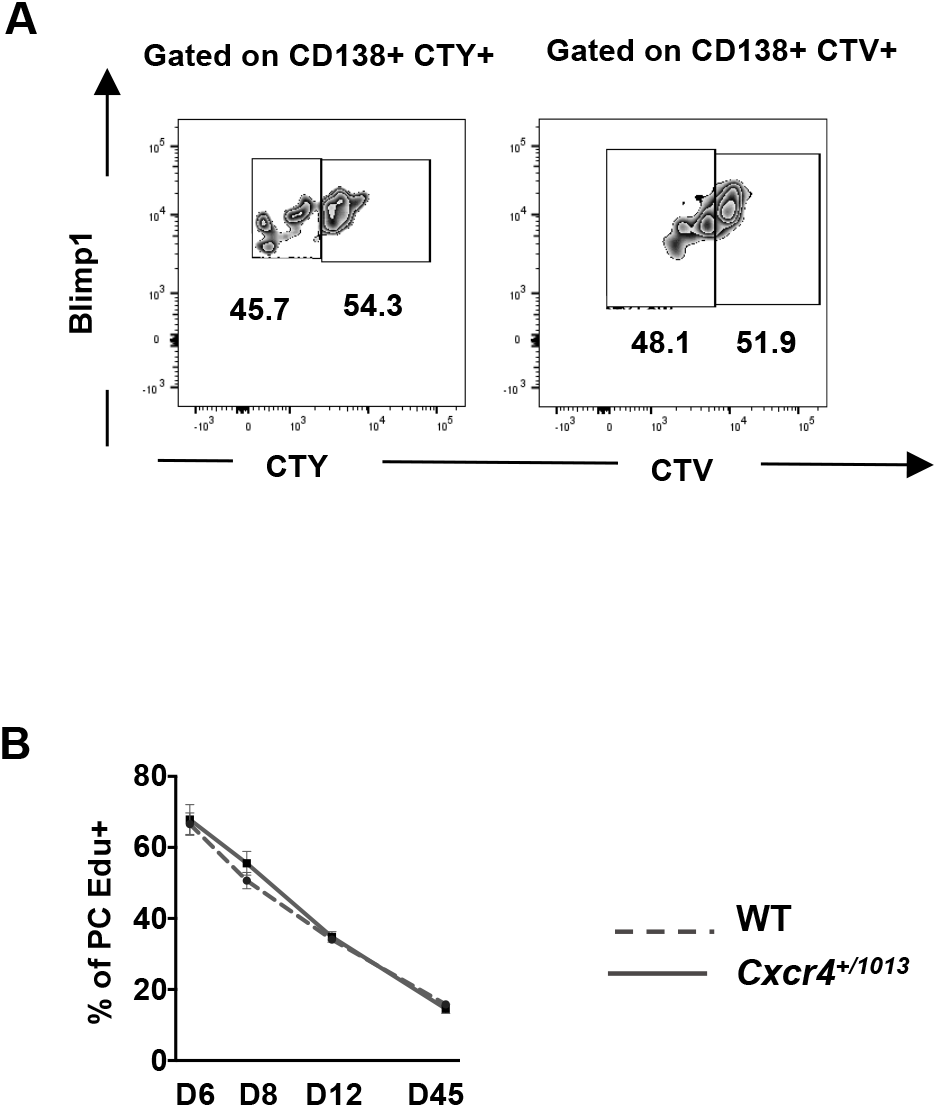
Cxcr4 desensitization do not modulate PB proliferation: **(A)** Representative dot plots showing the dilution and hence the proliferation of CTV+ (*Cxcr4*^+/*1013*^ x Blimp1^GFP/+^) and CTY+ (WT x Blimp1^GFP/+^) PB/PCs in the BM of recipient mice (no or few divisions = CT High; several divisions= CT Low) in both genotypes at day 2 post transfer. **(B)** Frequency of EdU+ PCs from d6 and during the chase at the indicated time-points after NP-LPS immunisation. (n=4-5)

**Table S1:** List of antibodies used in flow cytometry.

| **Antibody** | **Clone** | **Host/Isotype** | **Supplier** |
| --- | --- | --- | --- |
| Anti-CD138 | 281-2 | Rat IgG2a, $\kappa$ | BD Biosciences |
| Anti-CD45R/B220 | RA3-6B2 | Rat / IgG2a, kappa | BD Biosciences |
| Anti-CD19 | 1D3 | Rat IgG2a, $\kappa$ | BD Biosciences |
| Anti-CD21/35 | 7G6 | Rat gG2b, $\kappa$ | BD Biosciences |
| Anti-CD23 | B3B4 | Rat IgG2a, $\kappa$ | BD Biosciences |
| Anit-Vla-4 (CD49d) | R1-2 | Rat IgG2b, $\kappa$ | BD Biosciences |
| Anti-Lfa-1 (CD11a) | 2D7 | Rat IgG<sub>2a</sub>, $\kappa$ | BD Biosciences |
| Anti-CD44 | IM7 | Rat IgG2b, $\kappa$ | BD Biosciences |
| Anti-Ki67 | B56 | Mouse IgG<sub>1</sub>, $\kappa$ | BD Biosciences |
| Anti-Cxcr4 | 2B11 | Rat / IgG2b, kappa | BD Biosciences |
| Anti-Cxcr3 | HTK888 | Armenian Hamster IgG | BioLegend |

**Table S2:**
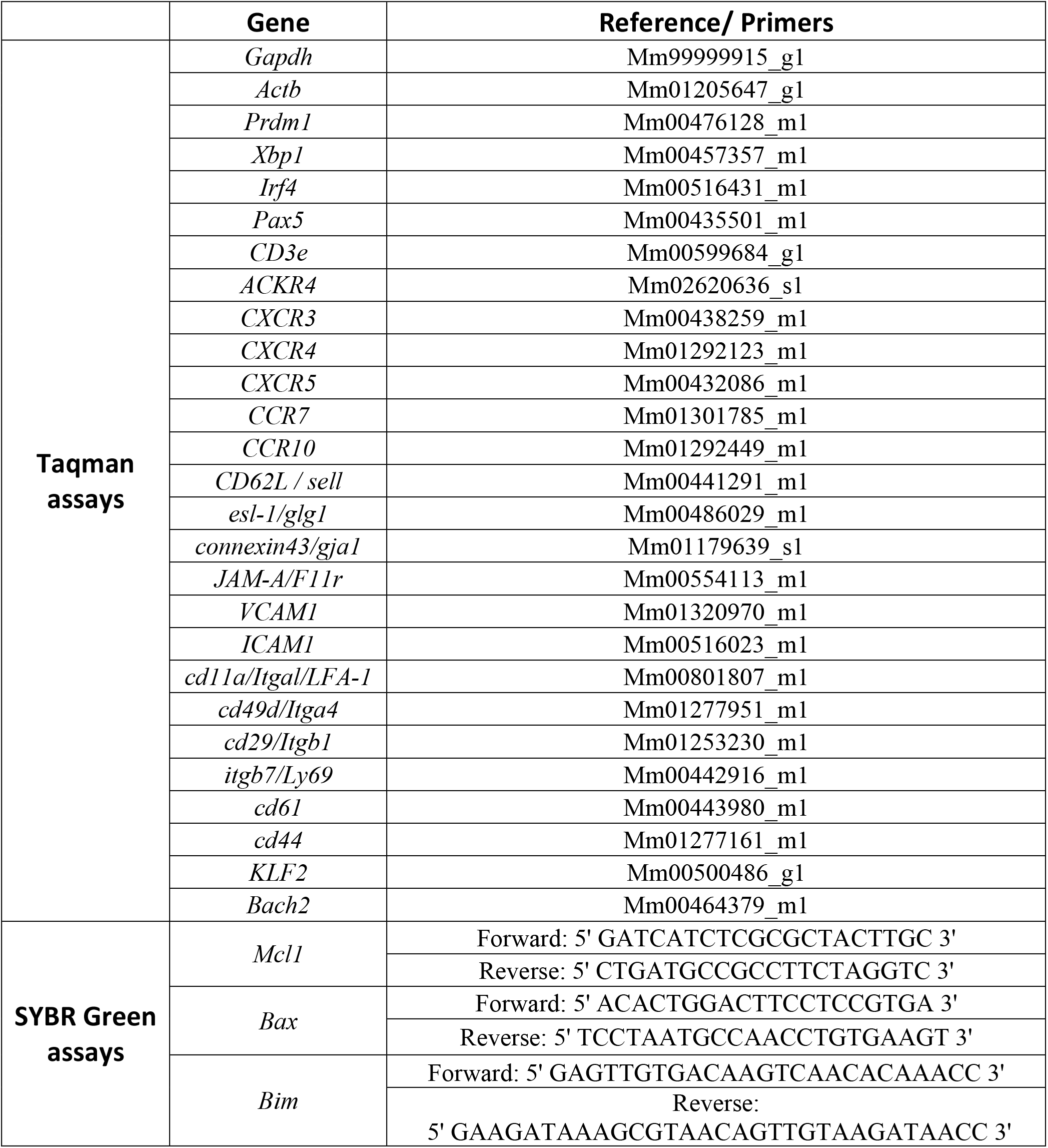
List of primers used for RT-qPCR.

## References

1. Nagasawa, T., Kikutani, H. & Kishimoto, T. Molecular cloning and structure of a pre-B-cell growth-stimulating factor. Proc. Natl. Acad. Sci. U. S. A. 91, 2305–2309 (1994).

2. Nagasawa, T. et al. Defects of B-cell lymphopoiesis and bone-marrow myelopoiesis in mice lacking the CXC chemokine PBSF/SDF-1. Nature 382, 635–8 (1996).

3. Bannard, O. et al. Germinal center centroblasts transition to a centrocyte phenotype according to a timed program and depend on the dark zone for effective selection. Immunity 39, 912–924 (2013).

4. Allen, C. D. C. et al. Germinal center dark and light zone organization is mediated by CXCR4 and CXCR5. Nat. Immunol. 5, 943–952 (2004).

5. Hargreaves, D. C. et al. A coordinated change in chemokine responsiveness guides plasma cell movements. J. Exp. Med. 194, 45–56 (2001).

6. Nie, Y. et al. The Role of CXCR4 in Maintaining Peripheral B Cell Compartments and Humoral Immunity. J. Exp. Med. 200, 1145–1156 (2004).

7. Biajoux, V. et al. Efficient Plasma Cell Differentiation and Trafficking Require Cxcr4 Desensitization. Cell Rep. 17, 193–205 (2016).

8. Victora, G. D. et al. Germinal center dynamics revealed by multiphoton microscopy with a photoactivatable fluorescent reporter. Cell 143, 592–605 (2010).

9. Moore, C. A. C., Milano, S. K. & Benovic, J. L. Regulation of Receptor Trafficking by GRKs and Arrestins. Annu. Rev. Physiol. 69, 451–482 (2007).

10. Balabanian, K. et al. WHIM syndromes with different genetic anomalies are accounted for by impaired CXCR4 desensitization to CXCL12. (2005). doi:10.1182/blood-2004

11. Kawai, T. et al. Enhanced function with decreased internalization of carboxy-terminus truncated CXCR4 responsible for WHIM syndrome. Experimental Hematology 33,(2005).

12. Gulino, A. V. et al. Altered leukocyte response to CXCL12 in patients with warts hypogammaglobulinemia, infections, myelokathexis (WHIM) syndrome. Blood 104, 444–452 (2004).

13. Balabanian, K. et al. Proper desensitization of CXCR4 is required for lymphocyte development and peripheral compartmentalization in mice. (2012). doi:10.1182/blood-2012

14. Treon, S. P. et al. Somatic mutations in MYD88 and CXCR4 are determinants of clinical presentation and overall survival in Waldenström macroglobulinemia. Blood 123, 2791–2796 (2014).

15. Poulain, S. et al. Genomic landscape of CXCR4 mutations in Waldenström macroglobulinemia. Clin. Cancer Res. 22, 1480–1488 (2016).

16. Treon, S. P., Xu, L. & Hunter, Z. MYD88 Mutations and Response to Ibrutinib in Waldenström’s Macroglobulinemia. N. Engl. J. Med. 373, 584–586 (2015).

17. Leblond, V. et al. Treatment recommendations from the Eighth International Workshop on Waldenström’s Macroglobulinemia. Blood 128, 1321–1328 (2016).

18. Fairfax, K. A., Kallies, A., Nutt, S. L. & Tarlinton, D. M. Plasma cell development: From B-cell subsets to long-term survival niches. Seminars in Immunology 20, 49–58 (2008).

19. Hasbold, J., Corcoran, L. M., Tarlinton, D. M., Tangye, S. G. & Hodgkin, P. D. Evidence from the generation of immunoglobulin G-secreting cells that stochastic mechanisms regulate lymphocyte differentiation. Nat. Immunol. (2004). doi:10.1038/ni1016

20. Oliver, A. M., Martin, F., Gartland, G. L., Carter, R. H. & Kearney, J. F. Marginal zone B cells exhibit unique activation, proliferative and immunoglobulin secretory responses. Eur. J. Immunol. 27, 2366–2374 (1997).

21. Peled, A., Klein, S., Beider, K., Burger, J. A. & Abraham, M. Role of CXCL12 and CXCR4 in the pathogenesis of hematological malignancies. Cytokine 109, 11–16 (2018).

22. Ngo, H. T. et al. SDF-1/CXCR4 and VLA-4 interaction regulates homing in Waldenstrom macroglobulinemia. Blood 112, 150–158 (2008).

23. Ngo, H. T. et al. SDF-1/CXCR4 and VLA-4 interaction regulates homing in Waldenstrom macroglobulinemia. (2008). doi:10.1182/blood-2007-12-129395

24. Hunter, Z. R. et al. Transcriptome sequencing reveals a profile that corresponds to genomic variants in Waldenström macroglobulinemia. Blood 128, 827–38 (2016).

25. Weisel, F. J., Zuccarino-Catania, G. V., Chikina, M. & Shlomchik, M. J. A Temporal Switch in the Germinal Center Determines Differential Output of Memory B and Plasma Cells. Immunity 44, 116–130 (2016).

26. Triantafilou, M. et al. Chemokine receptor 4 (CXCR4) is part of the lipopolysaccharide ‘sensing apparatus’. Eur. J. Immunol. 38, 192–203 (2008).

27. Triantafilou, K., Triantafilou, M. & Dedrick, R. L. A CD14-independent LPS receptor cluster. Nat. Immunol. 2, 338–345 (2001).

28. Kishore, S. P., Bungum, M. K., Platt, J. L. & Brunn, G. J. Selective suppression of Toll-like receptor 4 activation by chemokine receptor 4. FEBS Lett. 579, 699–704 (2005).

29. Freitas, C. et al. Lymphoid differentiation of hematopoietic stem cells requires efficient Cxcr4 desensitization. J. Exp. Med. 214, 2023–2040 (2017).

30. Barwick, B. G., Scharer, C. D., Bally, A. P. R. & Boss, J. M. Plasma cell differentiation is coupled to division-dependent DNA hypomethylation and gene regulation. Nat. Immunol. 17, 1216–1225 (2016).

31. Scharer, C. D., Barwick, B. G., Guo, M., Bally, A. P. R. & Boss, J. M. Plasma cell differentiation is controlled by multiple cell division-coupled epigenetic programs. Nat. Commun. 9, (2018).

32. Caron, G. et al. Cell-Cycle-Dependent Reconfiguration of the DNA Methylome during Terminal Differentiation of Human B Cells into Plasma Cells. Cell Rep. 13, 1059–71 (2015).

33. Hunter, Z. R. et al. The genomic landscape of Waldenström macroglobulinemia is characterized by highly recurring MYD88 and WHIM-like CXCR4 mutations, and small somatic deletions associated with B-cell lymphomagenesis. Blood 123, 1637–1646 (2014).

34. Cao, Y. et al. The WHIM-like CXCR4 S338X somatic mutation activates AKT and ERK, and promotes resistance to ibrutinib and other agents used in the treatment of Waldenstrom’s Macroglobulinemia. Leukemia 29, 169–176 (2015).

35. Roccaro, A. M. et al. C1013G/CXCR4 acts as a driver mutation of tumor progression and modulator of drug resistance in lymphoplasmacytic lymphoma. (2014). doi:10.1182/blood-2014-03-564583

36. Hui, L. & Chen, Y. Tumor microenvironment: Sanctuary of the devil. Cancer Lett. 368, 7–13 (2015).

37. Kallies, A. et al. Plasma Cell Ontogeny Defined by Quantitative Changes in Blimp-1 Expression. J. Exp. Med. J. Exp. Med 200, 967–977 (2004).

38. Kusumbe, A. P., Ramasamy, S. K., Starsichova, A. & Adams, R. H. Sample preparation for high-resolution 3D confocal imaging of mouse skeletal tissue. Nat. Protoc. 10, 1904–1914 (2015).

39. Thai, L. H. et al. BAFF and CD4+ T cells are major survival factors for long-lived splenic plasma cells in a B-cell–depletion context. Blood 131, 1545–1555 (2018).

